# Organization of an Ascending Circuit that Conveys Flight Motor State

**DOI:** 10.1101/2023.06.07.544074

**Authors:** Han S. J. Cheong, Kaitlyn N. Boone, Marryn M. Bennett, Farzaan Salman, Jacob D. Ralston, Kaleb Hatch, Raven F. Allen, Alec M. Phelps, Andrew P. Cook, Jasper S. Phelps, Mert Erginkaya, Wei-Chung A. Lee, Gwyneth M. Card, Kevin C. Daly, Andrew M. Dacks

## Abstract

Natural behaviors are a coordinated symphony of motor acts which drive self-induced or reafferent sensory activation. Single sensors only signal presence and magnitude of a sensory cue; they cannot disambiguate exafferent (externally-induced) from reafferent sources. Nevertheless, animals readily differentiate between these sources of sensory signals to make appropriate decisions and initiate adaptive behavioral outcomes. This is mediated by predictive motor signaling mechanisms, which emanate from motor control pathways to sensory processing pathways, but how predictive motor signaling circuits function at the cellular and synaptic level is poorly understood. We use a variety of techniques, including connectomics from both male and female electron microscopy volumes, transcriptomics, neuroanatomical, physiological and behavioral approaches to resolve the network architecture of two pairs of ascending histaminergic neurons (AHNs), which putatively provide predictive motor signals to several sensory and motor neuropil. Both AHN pairs receive input primarily from an overlapping population of descending neurons, many of which drive wing motor output. The two AHN pairs target almost exclusively non-overlapping downstream neural networks including those that process visual, auditory and mechanosensory information as well as networks coordinating wing, haltere, and leg motor output. These results support the conclusion that the AHN pairs multi-task, integrating a large amount of common input, then tile their output in the brain, providing predictive motor signals to non-overlapping sensory networks affecting motor control both directly and indirectly.

## Introduction

Animals exploit a multisensory strategy to navigate their environment. In doing so, the animal’s own movements can activate one or more sensory modes in the process of reafference, that must be reliably distinguished from sensory activation from outside stimuli (exafference) to coordinate behavior [1]. However, individual sensory organs or structures only signal the presence and magnitude of sensory cues, but cannot provide source information. For instance, a mechanosensory hair will activate when bent by an external object, such as a predator, or by the animal’s own movement, such as when brushing up against a stationary object. To distinguish reafference from exafference, the central nervous system implements a broad class of feedforward circuits, commonly referred to as corollary discharge circuits (CDCs), which provide predictive motor information to sensory and motor pathways [2]. CDCs impact sensorimotor integration via diverse means including modulation of network processing [3]; [4], blanket suppression or temporally precise inhibitory gating of sensory processing [5–8], and modulating efferent pathways that tune sensory sensitivity [9,10]. Importantly, CDCs are often modulatory in nature and can up or downregulate responsiveness of a sensory neuropil to reafferent signals [4,10]. Throughout the animal kingdom, CDCs convey information from a variety of motor control centers to most, if not all, sensory modalities. The fundamental importance of these predictive motor signals is highlighted by their failure, which results in attributional errors associated with nearly every form of sensory hallucination whether fatigue-[11,12] or disease-induced [13,14]. Thus, predictive motor signals from CDCs are essential for the animal to effectively use sensory information to make adaptive choices that optimize behavioral performance.

Despite being studied in diverse species and sensory modalities, there remain many open questions about the cellular and synaptic mechanisms underlying CDC function. For example, a precise efference copy of motor commands can be physiologically observed during visual processing in *Drosophila* [8], but the cellular basis of this signal has not yet been established. Furthermore, CDCs representing different motor information can converge onto a given sensory pathway [6,9,15–17], yet the combined impact of this convergence remains unknown. Finally, a single CDC can also distribute information to multiple sensory and/or motor neuropils [4,7], yet functional consequences of this distribution remain unknown.

We recently described two pairs of histaminergic neurons, which originate within the ventral nerve cord (VNC) and project to the brain of the moth *Manduca sexta [18]*. These ascending histaminergic neurons (AHNs) have somas in the mesothoracic (MsAHNs) and metathoracic (MtAHNs) neuromeres. Across insect taxa, the AHNs bilaterally ramify the subesophageal zone (SEZ) and the antennal mechanosensory and motor center (AMMC). In night-flying plume tracking insects, the MsAHNs have also been co-opted into the antennal lobe [19] and in *Manduca*, only a small number of GABAergic local interneurons (LNs) express the inhibitory histamine B receptor suggesting that the AHNs affect local processing of odor information through a disinhibitory network mechanism [18]. Paired recordings of the MsAHNs and a primary wing motor fiber indicates that MsAHN firing rate increases prior to wing motor output from the VNC, thus, their activation is thought to be the consequence of receiving direct descending wing motor command signals. AHN activation therefore results in disinhibition within the antennal lobe just in advance of flight, resulting in an upregulation of temporal precision with which antennal lobe projection neurons entrain to the stimulus temporal structure [4], such as that induced by the beating wings [20,21]. The consequence of this increased temporal fidelity is associated with enhanced sensory acuity [4]. Thus, in *Manduca* the MsAHNs represent a CDC that informs the AL of wing motor action, allowing it to upregulate olfactory processing and performance during flight. However, the cellular and synaptic mechanisms that mediate AHN activity, as well as the functional role of the AHNs across the other projection zones remain unexplored.

To resolve the broader network architecture within which the AHNs are integrated, and the mechanisms by which they affect sensorimotor integration, we turned to the wealth of approaches afforded by *Drosophila melanogaster*. We asked to what extent are the two AHN pairs integrated within the same sensorimotor networks? Furthermore, do the AHNs represent motor information in this model and if so, are they activated under the same behavioral contexts? Finally, do the AHN pairs follow the same organizational principles with respect to their synaptic connectivity and mechanisms of communication, or do they represent operationally different circuits despite their similarities in basic morphology and transmitter content? To address these questions, we exploited connectomics, molecular, anatomical, physiological and behavioral approaches to comprehensively map synaptic connectivity and explore the function of the AHNs.

## Results

### AHN Anatomical Characterization Throughout the CNS

Histaminergic neurons project throughout the brain and VNC of *Drosophila* (Figure 1A; [22–26] and innervate most neuropil. Similar to *Manduca* and many other insect species [19], there are two pairs of ascending histaminergic neurons (AHNs) with somas located within the mesothoracic (the MsAHNs) and metathoracic (the MtAHNs) neuromeres. The remaining histaminergic soma in the VNC reside in the abdominal segment and do not ascend through the neck connective (Figure 1A). We therefore first aimed to determine the contribution of the AHNs to the total histaminergic projections observed within the central nervous system. To this end, we identified several driver lines that include the AHNs, but not other histaminergic neurons (Figure S1A-E). Using one such line, we drove the expression of diphtheria toxin A [27] to ablate the AHNs. This approach eliminated histamine labeling in the thoracic neuromeres (Figure 1B) as well as in several brain neuropils, including most of the sub-esophageal zone (SEZ), antennal mechanosensory and motor center (AMMC) (Figure 1B’), the saddle and the posterior slope (Figure 1B”). To resolve the independent projections of the AHNs, we used the MultiColor FLP Out approach (MCFO; [28] to stochastically label single AHNs (Figure 1C & D). This established that individual MsAHNs arborize within both the ipsilateral and contralateral meso and pro-thoracic neuromeres before ascending to the brain to innervate the SEZ, saddle and posterior slope (Figure 1C’-C’’), whereas single MtAHNs unilaterally arborize in the GNG, AMMC (Figure 1D) and all leg neuropils contralateral to the soma (Figure 1D’).

**Figure 1.**
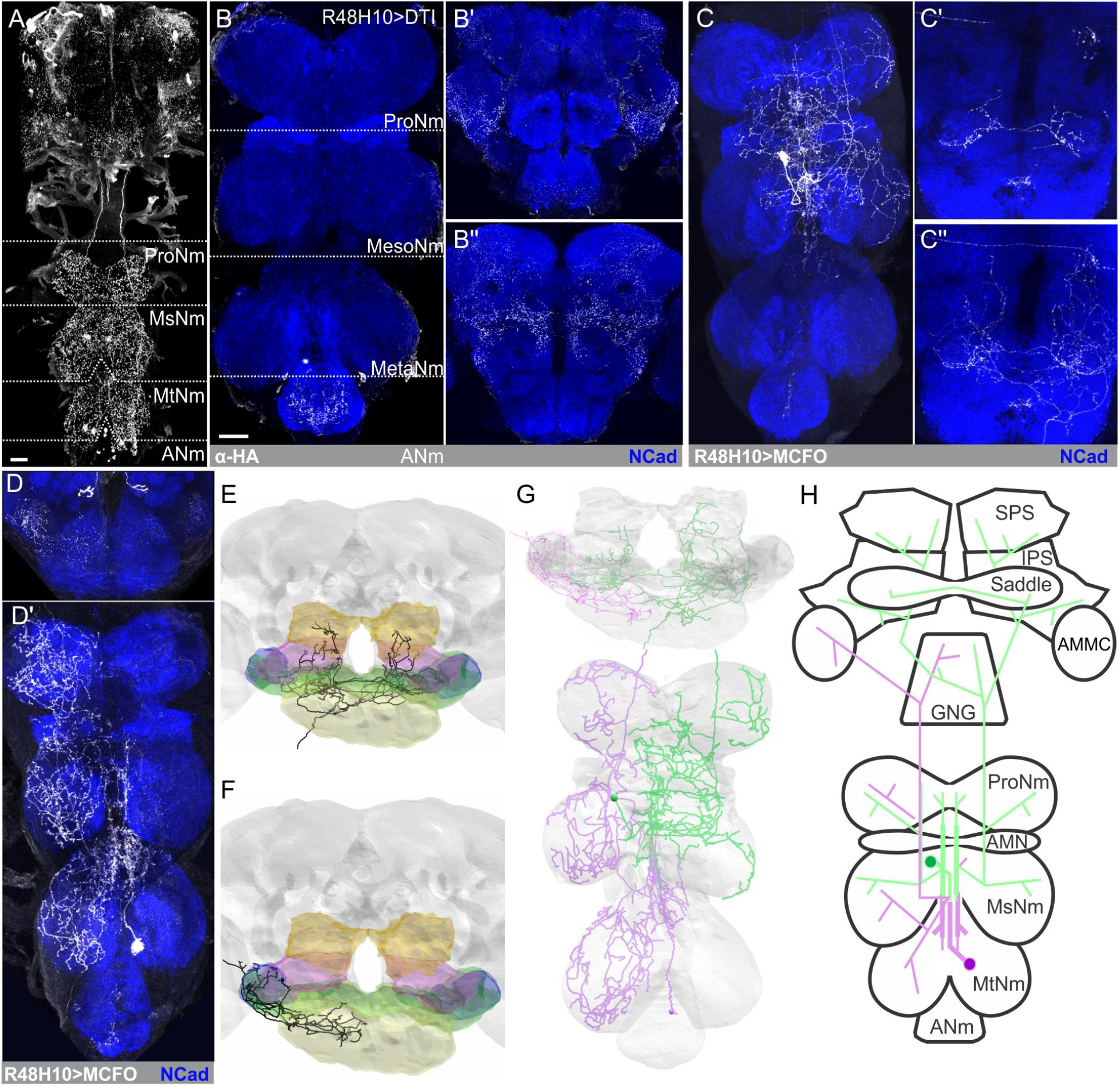
General AHN morphology and histamine expression in the CNS. **A)** Histamine immunolabeling in the intact CNS of Drosophila melanogaster. ProNM; prothoracic neuromere, MsNM; Mesothoracic neuromere, MtNM; Metathoracic neuromere, ANM; Abdominal neuromere. **B)** Histamine immunolabeling in R48H10-Gal4 flies driving expression of diptheria toxin-A in **B)** the VNC as well as **B’)** anterior and **B’’)** posterior depths in the brain. **C)** Single MsAHN clone within the **C)** VNC and at **C’)** medium and **C’’)** posterior depths of the brain. **D)** Single MtAHN clone within the **D)** VNC and at **D’)** anterior depths of the brain. **E, F)** Reconstruction of a **E)** MsAHN and **F)** MtAHN within the Female Adult Fly Brain (FAFB) EM volume. Gnathal ganglion (GNG; yellow), saddle (green), antennal mechanosensory motor center (AMMC; blue), inferior posterior slope (IPS; magenta), superior posterior slope (SPS; orange). **G)** Reconstruction of a MtAHN (lavender) and MsAHN (green) in the FAFB and Female Adult Nerve Cord EM volumes. Scale bars = 20μm. **H)** Cartoon schematics of the MtAHN (lavender) and MsAHN (green).

Using their unique morphology, we then located candidate MsAHNs and MtAHNs within three large electron microscopy volumes, the Female Adult Fly Brain (FAFB; [29], the Female Adult Nerve Cord (FANC; [30,31] and the Male Adult Nerve Cord (MANC; [32,33] datasets (Figure 1D, F & G). In the brain, we identified the MsAHNs based on their expected bilateral arborization patterns in the SEZ, saddle and posterior slope (Figure 1E) and the MtAHNs based on their unilateral innervation of the AMMC and dorsal GNG (Figure 1F). Furthermore, we identified the MsAHNs and MtAHNs within the male and female VNC datasets, based on soma position and morphology, then reconstructed them (Figure 1G). The general morphology of the AHNs was similar between male and female datasets (Figure S1F-I and K), except for additional processes from the MsAHNs projecting laterally and ventrally within the prothoracic neuromeres in FANC that were absent in MANC (Figure S1H-I). However, we observed variability for the ventrolaterally projecting prothoracic branch among single MCFO clones of MsAHNs from male flies, suggesting that these morphological differences represent individual, rather than sex-specific differences (Figure S1J). There were no obvious differences in the morphology of the MtAHNs between the FANC and MANC datasets (Figure S1K). Thus, MCFO-derived single AHNs (Figure 1C-D) and EM-based reconstructions indicate that the AHNs pairs partially overlap within the VNC but are non-overlapping in the brain (Figure 1H). To determine whether other neurons represented good AHN candidates, we took two approaches. First, we manually traced the major processes of all neurons within the same ventral-to-dorsal tracts through which the MsAHNs and MtAHNs soma project within the FANC dataset, and found no neurons whose coarse morphology were consistent with the features of the AHNs, as revealed by MCFO (Figure 1C-D). Second, we used NBLAST [34] with our AHN candidates to search for similar cells within the MANC volume (Figure S1L-M), but again none of the top hits were similar to the single AHN neurons observed with light microscopy (Figure 1C-D). Thus, both methods suggest that the AHN candidate cells found in MANC and FANC are high-confidence matches to AHNs.

Next, we characterized primary input and output regions of the AHNs using driver lines that restricted Gal4 expression exclusively to either the MsAHNs or MtAHNs. Using the somatic/dendritic marker DenMark (ICAM5-RFP) and the presynaptic marker synaptotagmin tagged with GFP (syteGFP) [35], we labeled the primary input and output regions of the AHNs, and compared these immunohistochemical results with EM volume reconstructions where we annotated pre- and postsynaptic sites (Figure 2). Within the VNC, the MsAHNs had sparse expression of syteGFP throughout the prothoracic and mesothoracic neuromeres (Figure 2A) and prominent ICAM5-RFP labeling primarily localized in the wing and intermediate tectulum (Figure 2A & B), [36]; [37]. Within the brain, only syteGFP was present in the SEZ, saddle and posterior slope (Figure 2C & D), consistent with FAFB synapse annotations which are predominantly presynaptic within the brain (Table 4). The distribution of manually annotated postsynaptic sites of the MsAHN EM reconstructions were highly consistent with the transgenic expression of ICAM5-RFP by the MsAHN splitGal4 line, but relatively more presynaptic sites within leg neuropil were evident in the MsAHNs within the FANC EM volume (Figure 2E). The MtAHNs on the other hand had syteGFP expression in all three thoracic neuromeres (Figure 2F & G), as well as the AMMC (Figure 2H) and dorsal GNG (Figure 2I). MtAHN expression of ICAM5-RFP was distributed throughout the posterior intermediate tectulum of the metathoracic neuromere, and extended into the mesothoracic neuromere (Figure 2F & G) collectively spanning neuropil involved in leg and wing coordination [36,37]. As with the MsAHNs, the MtAHN transgenic labeling of pre- and postsynaptic zones was consistent with EM volume reconstruction’s distribution of pre- and postsynaptic sites (Figure 2E & 2J). Taken together, these results provide strong confirmation that we have identified the AHNs in the FANC and FAFB connectomic volumes. Importantly, the AHNs receive the bulk of their input within the medial regions of the VNC associated with locomotor control, and project local output extensively within the leg and wing sensory-motor neuropil and multiple sensory processing regions in the brain.

**Figure 2.**
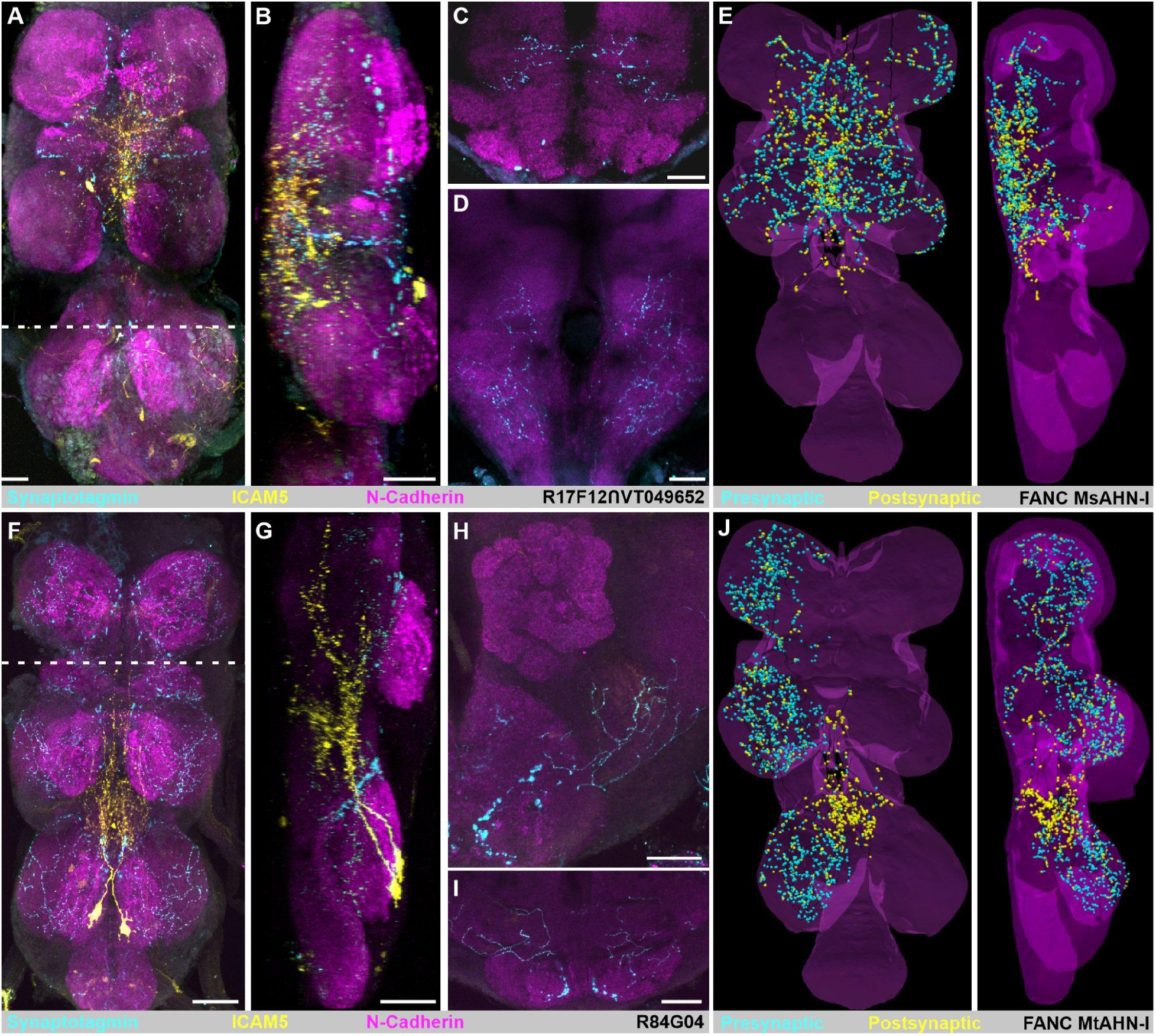
Distribution of AHN input and output regions. **A)** Input and output regions of the MsAHNs. R17F12 ∩ VT049652-Gal4 driving expression of the axon terminal marker synaptotagmin-eGFP (“syteGFP”; cyan) and the dendrite/soma marker ICAM5-mCherry (“DenMark”; yellow). NCAD serves as a neuropil marker (magenta). **A)** Horizontal view of MsAHN syteGFP and DenMark expression within the VNC. Dashed line indicates the border of the sagittal view in **B**. **B)** Sagittal view of image stack in **A)**. **C)** Frontal view of MsAHN syteGFP and DenMark expression within the saddle. **D)** Frontal view of MsAHN syteGFP and DenMark expression within the posterior slope. **H)** Reconstruction of the MsAHN from the FANC EM volume with presynaptic (cyan) and postsynaptic (yellow) marked. **F)** Input and output regions of the MtAHNs. R84G04-Gal4 driving expression of the axon terminal marker synaptotagmin-eGFP (“syteGFP”; cyan) and the dendrite/soma marker ICAM5-mCherry (“DenMark”; yellow). NCAD serves as a neuropil marker (magenta). Dashed line indicates the border of the sagittal view in **G**. **G)** Sagittal view of image stack in **F)**. **H)** Frontal view of MtAHN syteGFP and DenMark expression within the AMMC. **I)** Frontal view of MtAHN syteGFP and DenMark expression within the GNG. **J)** Reconstruction of the MtAHN from the FANC EM volume with presynaptic (cyan) and postsynaptic (yellow) marked. Scale bars = 20μm.

Next, we used the FANC and MANC EM datasets to retrieve or reconstruct the upstream synaptic partners of the AHNs to determine the possible information sources integrated by the AHNs. We first classified synaptic partners into one of 6 broad neuronal classes; sensory, ascending sensory, ascending (central), descending, interneurons (restricted to the VNC), or motor neurons. All four AHNs had relatively similar proportional demographics (across FANC and MANC), with descending neurons representing the majority of input to all AHNs (50-68%) followed by interneurons (17-37%), with relatively small contributions from the other cell class (Figure 3A & B). To determine if different upstream partner neuronal classes synapse upon distinct regions of the AHNs, we plotted the location of input sites to the AHNs based on the broad neuronal class of the upstream partner (Figure 3C, D, Figure S2A-H). While interneurons synapse uniformly across the AHN skeletons (Figure S2B, F), the descending (Figure S2A, E) and ascending neurons (Figure S2C, G) primarily synapse upon medial regions of the AHNs near the midline, in particular within tectular neuropils. Sensory neurons tended to synapse upon the more proximal branches, and for the MtAHNs made contact almost exclusively within the mesothoracic leg neuropil (Figure S2D, H).

**Figure 3.**
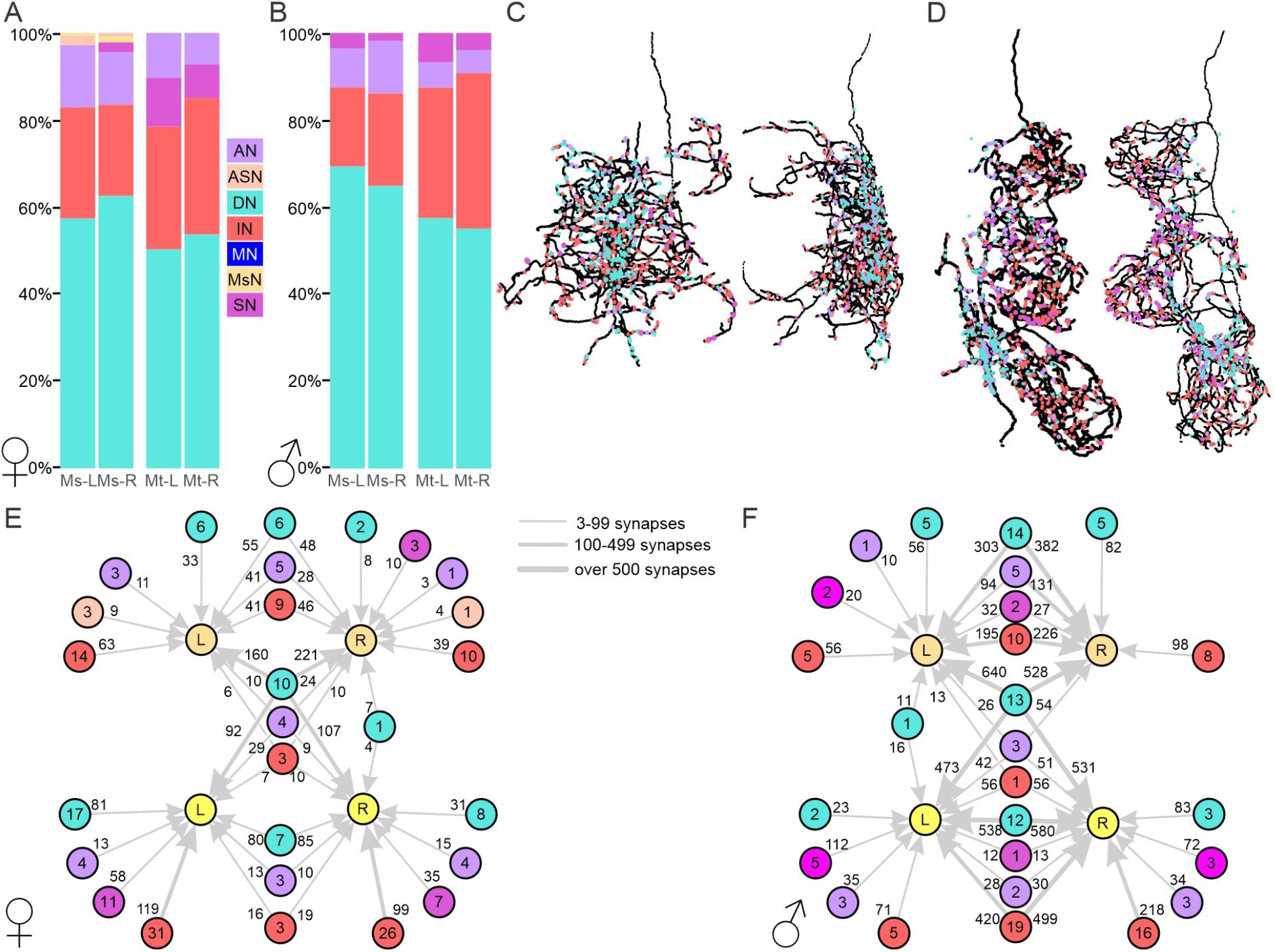
Connectivity of upstream inputs to the AHNs. **A-B)** Synapse fractions expressed as percent of the total among identified upstream partners for the 6 classes of neuron type in **A)** FANC and **B)** MANC. **C)** Horizontal (left) and sagittal (right) views of the synapse distributions upon the left MsAHN in FANC from descending neurons (cyan), interneurons (red), ascending neurons (lavender) and sensory neurons (pink). **D)** Horizontal (left) and sagittal (right) views of the synapse distributions upon the left MtAHN in FANC from descending neurons (cyan), interneurons (red), ascending neurons (lavender) and sensory neurons (pink). **E-F)** Graph plot of upstream partners to the AHNs in the **E)** FANC and **F)** MANC dataset. Node number indicates number of neurons within a neuronal category. Edge number indicates total number of synaptic connections. Abbreviations: ascending neuron (AN), ascending sensory neuron (ASN), descending neuron (DN), interneuron (IN), motor neuron (MN), MsAHN (MsN), sensory neuron (SN).

Depending upon the degree of convergence of the upstream partners on all four AHNs, the AHNs could represent common information or information unique to single cells or pairs of AHNs. To determine the degree to which individual AHNs receive common versus distinctive input, we generated graph plots to depict the input to each AHN by cell class (Figure 3E-F). Among the five cell classes that provide input to the AHNs, all five classes provide unique input to the MsAHNs and four provide unique input to the MtAHNs. Four classes provide bilateral input to either the MsAHNs or the MtAHNs. However, the primary common input to all four AHNs was contributed almost exclusively by DNs (Figure 3E-F), with the exception of three ascending and one interneuron in FANC and two ascending neurons in MANC. Finally, the number of synapses from both common inputting DNs and bilaterally inputting DNs was substantial relative to the other cell classes. This implies that while there may be diverse input to individual or pairs of AHNs, the “unifying” source of input to the AHNs derives from descending signals. Thus, the AHN pairs likely integrate common patterns of descending command signals.

A large catalog of DNs in *Drosophila* have been classified at the light-level by morphology and functional organization in the CNS [36,38], and most of these DNs are now further identified in the MANC EM volume [39]. Guided by these works, we identified in both FANC and MANC specific DN types that provide substantial input to single, pairs, or all four AHNs (Figure 4A & B). Although the AHN pairs receive input from a unique combination of DNs, nearly 40% of DNs in MANC (46% of total DN synapses) converge onto both AHN pairs (Figure 4C-E, Figure S3A), in particular from the DNg02s (Figure 4F-G) and DNp54s. While little is known about the DNp54s, the DNg02s represent flight steering command neurons that are responsive to visual motion during flight and whose activation increases wing stroke amplitude [38], to adjust flight path during the optomotor response of flies [40]. Using a reporter of choline acetyltransferase translation [41], we determined that the majority of DNg02s are cholinergic (Figure 4H), but express neither the vesicular glutamate transporter or GABA (Figure S3B, C) and thus likely provide excitatory input to the AHNs. The DNp54s did not co-express reporters for any small classical transmitters (Figure S3D-H). To test if the AHNs are activated by wing motor output, we carried out calcium imaging of the DNg02 population using jGCaMP7b and each AHN pair using jGCaMP7f during the induction of flight. Flies were dissected and imaged from the posterior side of the head for DNg02 dendrites or from the ventral side of the thorax for AHN soma, and a brief air puff was used to induce flight. Consistent with Namiki et al (2021), the DNg02s became active shortly after the induction of flight (Figure 4I). Likewise, both the MsAHNs (Figure 4J) and MtAHNs (Figure 4K) each became active after the induction of flight, while no change in fluorescence was observed when GFP was expressed in either AHN pair (Figure S3I, J), indicating that flight-induced changes in GCaMP signal were not movement artifacts. Flight-induced activation of AHNs suggests that they receive input correlated to wing movement, consistent with a predictive motor circuit. To test if the DNg02s provide excitatory, cholinergic drive to the AHNs, we expressed the light-activated ion channel CsChrimson [42] in the DNg02s while imaging from the MtAHN somata using jGCaMP7f in flies dissected from the ventral side of the thorax. Red-light activation of the DNg02s reliably resulted in a strong Ca^2+^ transient from the MtAHNs (Figure 4L), while red-light did not induce MtAHN Ca^2+^ transients when CsChrimson expression is driven by an “empty-Gal4” (a Gal4 line that doesn’t drive expression in the central nervous system; Figure S3M). Thus, the AHNs are activated during flight at least in part via excitatory input from the DNg02s. However, this does not preclude the possibility that other wing movements associated with courtship and aggression might also activate the AHNs.

**Figure 4.**
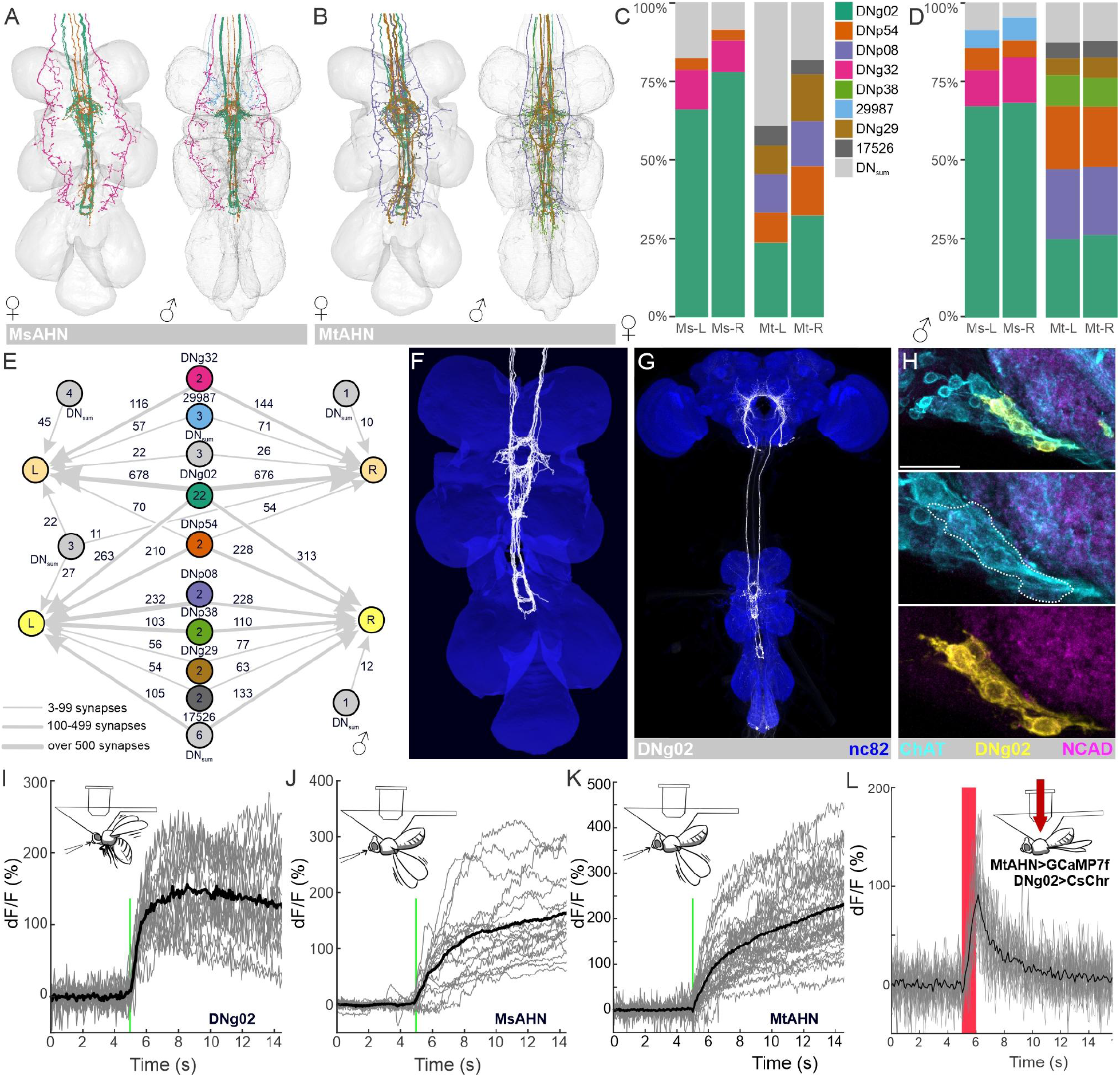
Connectivity of DNs upstream of the AHNs. **A-B)** Reconstruction of DNs representing greater than 5% of synaptic input from DNs to the **A)** MsAHNs and **B)** MtAHNs in both the FANC and MANC EM datasets. **C-D**) Synapse fractions of DNs upstream to the MsAHNs and MtAHNs in the **C**) FANC and **D**) MANC EM datasets. DNs are placed in their own category if the DN type has >5% connectivity with any AHN type, otherwise they are grouped as “DN_sum_”. The color scheme in the synapse fractions is matched to that of the reconstructions in **A** and **B**. **E)** Graph plot of DNs upstream to the MsAHNs and MtAHNs. Node number indicates number of neurons within a DN group. Edge number indicates total number of synaptic connections. Only connectivity from DNs to the AHNs is depicted. **F)** Reconstruction of the DNg02s (the largest source of synaptic input to the AHNs) in the FANC EM data set. **G)** VT039465-p65ADZ; VT023750-ZpGdbd (SS02625) splitGal4 line expressed in DNg02s (white). Nc82 used to delineate neuropil. Image courtesy of Shigehiro Namiki (cite Namiki 2018). **H)** Intersection between the DNg02 splitGal4 (yellow) and a ChAT-T2A-LexA (cyan) driver lines reveals that the DNg02s are cholinergic. NCAD (magenta) delineates neuropil and scale bar = 20μm. **I-K**) Flight-induced changes in Ca^2+^ levels measured via GCaMP7 expression in **I)** the DNg02s (6 flies, 3 trials), **J**) the MsAHNs (6 flies, 6 soma, 3 trials) and **K)** the MtAHNs (9 flies, 13 soma, 3 trials). Cartoon depicts orientation of flies during each recording and green line indicates timing of an air puff to trigger flight. Gray traces represent recordings from individual flies (DNg02) or soma (AHNs) and black trace represents the average Ca^2+^ transient across all animals. **L**) Ca^2+^ transients evoked measured via GCaMP7 expression in the MtAHNs in response to CsChrimson activation of the DNg02s. Gray traces represent recordings from individual AHN soma and black trace represents the mean fluorescence transient across all animals.

Based on the diversity of neurons that synapse upon the AHNs seen in the FANC and MANC datasets, we next sought to determine if the AHN pairs are subject to the same signaling molecules and second messenger pathways. In theory, the two pairs could receive synaptic input from the same upstream neurons, yet differ in their response to a given transmitter. By searching a single cell RNAseq library from the VNCs of five adult *Drosophila [43]* for histidine decarboxylase expression and then narrowing our search to those cells that express Hox genes associated with either the mesothoracic (*Antp*) and metathoracic (*Ubx*) neuromeres, we identified 9 candidate AHNs (5 MsAHNs and 4 MtAHNs) for which transcriptomic data was available. We then queried for expression of mRNA for known ionotropic and metabotropic receptors, G protein and second messenger effector protein subtypes, vesicular fusion proteins and ion channels (Figure 5, Figure S4A-C). Overall gene expression was relatively similar between the MsAHN and MtAHN candidates although they did cluster by cell pair when considering ion channels gene expression (Figure 5). Both AHN pairs expressed ionotropic receptors for the three primary transmitters (including nAChRα6, Rdl and GluClα) as well several GPCRs, most consistently Dop2R and DopEcR (Figure S4A). In particular, the nAChRα6 receptor (likely the mechanism of excitation through which the DNg02s act), the GABAA receptor (Rdl) and the Dop2R (Figure 5A) had relatively high expression levels in both AHN pairs. Three candidate MtAHNs clustered together when considering the expression patterns of genes associated with second messenger signaling (Figure S4B) including G-proteins themselves (such as Gαo, Gβ13F and Gɣ1), effector proteins (such as CamKII and PKA) and several genes involved in terminating signaling including Gprk2 [44] and dnc [45]. Thus, although there were some differences in neuronal gene expression between candidate AHN pairs, their overall expression was relatively similar.

**Figure 5.**
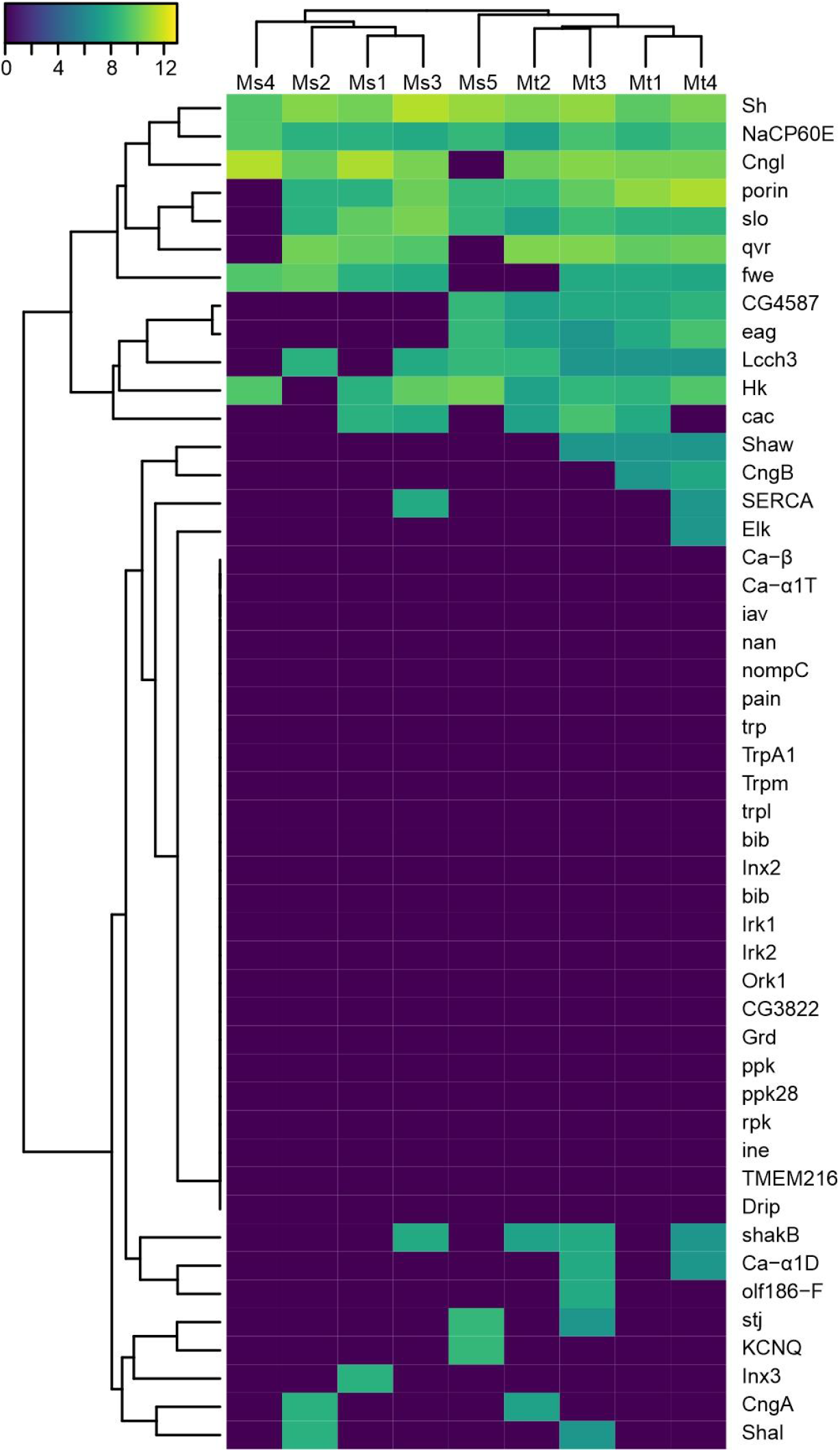
AHN ion channel expression profiling. Hierarchical cluster analysis showing normalized expression of ion channel genes across *Hdc*-expressing cells and either *Antp* (presumed MsAHN) or *Ubx* (presumed MtAHN). Values are read counts of each gene normalized by total counts per million (CPM) per cell, then log scaled (log2(n+1)).

We have shown that each AHN pair provides the sole source of histamine to several brain neuropils and collectively represent the sole source of histamine within the three thoracic neuromeres, where they possess partially overlapping output regions. Although both AHN types receive a portion of common input, the degree to which the AHN pairs converge upon common downstream partners remains unclear. The processes of the AHN pairs have the least overlap within the brain, suggesting the AHNs provide selective as opposed to convergent output. To confirm this, we traced 50% of downstream connections for a single MsAHN and MtAHN in the FAFB EM volume. Even without applying a synapse count threshold for inclusion, we observed virtually no common downstream synaptic partners of the AHN pairs (Figure 6A) within the brain, indicating that the different AHN types largely tile their output within the brain. We next examined the functional organization of the MsAHN output within its target brain neuropils (Figure 6B). The vast majority of the MsAHN output was directed to interneurons and a smaller proportion to descending neurons (Figure 6C). These downstream targets also receive input from lobula plate tangential cells (LPTCs; Figure 6D), which process visual motion [46–48]. Furthermore, brain interneurons targeted by MsAHNs provide synaptic input to the Giant fiber neuron, which trigger visually evoked escape behavior [49–53], and the DNa02 descending neuron (Figure 6D, Figure S5A) which modulates visually guided walking behavior [54]. The synaptic connectivity strengths of MsAHN downstream partners with LPTCs and DNa02 shown here are likely underestimated, as their connectivity was stochastically traced. Based on MsAHN projections into the posterior slope, a region associated with visual processing [55–57], and the connectivity of the MsAHNs to these descending neurons, it is possible that the MsAHNs modulate visually guided behavior within the context of ongoing motor output.

**Figure 6.**
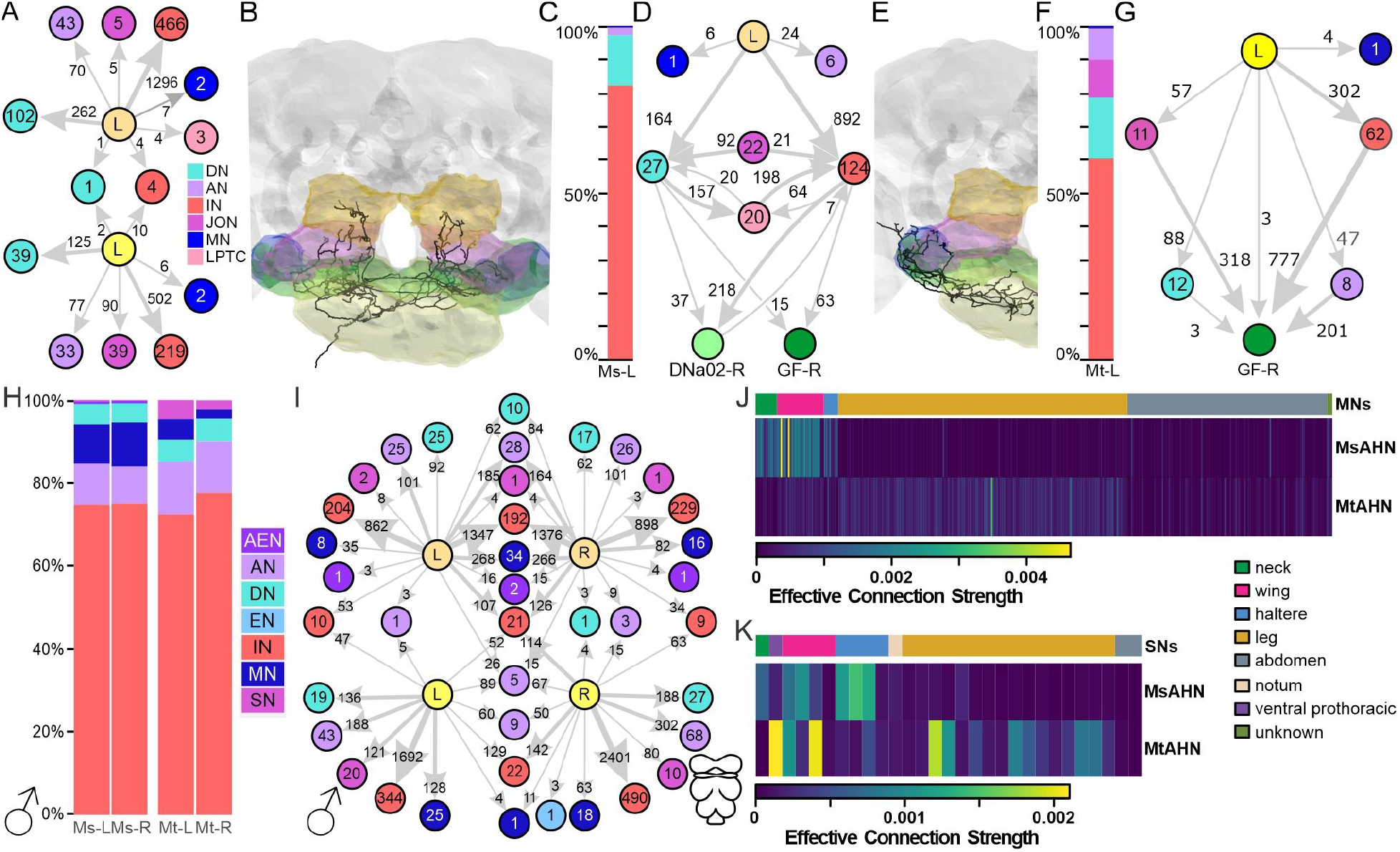
Downstream connectivity of the AHNs in the brain and female VN. **A)** Graph plot of the downstream targets of the left MsAHN (orange) and left MtAHN (yellow) by cell class. Ascending neurons (AN; lavender), descending neurons (DN; cyan), interneurons (IN; red), Johnston’s organ neurons (pink), lobula plate tangential cells (LPTC; salmon), motor neurons (MN; blue), reveals virtually no overlap in downstream targets, with no synapse threshold applied. **B)** Reconstruction of the left MsAHN in the FAFB dataset. Volumes for the gnathal ganglion (yellow), antennal mechanosensory and motor center (blue), saddle (green), inferior posterior slope (magenta) and superior posterior slope (orange) are highlighted. **C)** Synapse fraction plot of downstream partners of the left MsAHN within FAFB. Broad categories match the color scheme in **A)**. **D)** Graph plot of the downstream targets of the left MsAHN in FAFB. Threshold for inclusion was 3 synapses from the MsAHN onto each target. The MsAHN converged upon many common targets of the LPTCs including interneurons that provide input to two well-studied DNs, the giant fiber neuron (dark green) and DNa02 (light green). **E)** Reconstruction of the left MtAHN in the FAFB dataset. **F)** Synapse fraction plot of downstream partners of the MtAHN within FAFB. Broad categories match the color scheme in **A)**. **G)** Graph plot of the downstream targets of the left MtAHN in FAFB, and their connectivity with the giant fiber neuron (dark green). Threshold for inclusion was 3 synapses from the MtAHN onto each target. **H)** Synapse fraction of downstream partners of the MsAHNs and MtAHNs within the VNC of the MANC dataset. Threshold for inclusion was 3 synapses. Broad categories match the color scheme in **A)** with the addition of ascending efferent neurons (AEN; dark purple) which are those neurons with a soma within the VNC that project along peripheral nerves and the neck connective. **I)** Graph plot for the downstream partners of the MsAHNs (peach) and MtAHNs (yellow) from the MANC dataset. Threshold for inclusion was 3 synapses. **J)** Matrix of effective connection strength for connectivity between each AHN pair to individual motor neurons via one interneuron. Ordering of the matrix is based on the appendage or body part innervated by each motor neuron; wing (pink), leg (gold), neck (green), haltere (blue) or abdominal (grey). **K)** Matrix of effective connection strength for connectivity between each AHN pair to individual sensory neurons via one interneuron. Ordering of the matrix is based on the appendage or body part innervated by each sensory neuron. Sensory neurons are color coded as in **J)** with the addition of the notum (pink), ventral prothorax and unknown (dark green).

Within the brain, each MtAHN projects to the dorsal GNG and the AMMC (Figure 6E). The AMMC receives primary input from the Johnston’s organ (JO) at the base of the antennal aristae, which are tuned to different frequencies of acoustic and mechanosensory vibrations, as well as static deflections induced by forces such as gravity and wind [58–62]. Information transduced by the JO neurons is then distributed throughout diverse sensory and motor control networks found in regions such as the saddle, vest, wedge, and AMMC [63–68]. The MtAHN provided more input to more diverse cell classes relative to the MsAHN (Figure 6F). Although the majority of MtAHN synaptic output was directed towards interneurons, ∼15% was directed towards descending neurons and ∼10% each towards ascending and sensory neurons. Of the sensory afferents targeted by the MtAHNs, all were either JO-A or B type neurons which are sensitive to acoustic stimuli, and the majority of the remaining downstream targets of the MtAHNs in the brain were synaptically connected to the JO-A or B sensory afferents, implying a role for the MtAHNs in modulating or suppressing acoustic processing (Figure S5B). Interestingly, there was a high degree of convergence of all downstream cell classes onto the Giant Fiber neurons (Figure 6G, Figure S5C). In addition to responding to looming visual input, the Giant Fiber neurons also receive excitatory synaptic drive in response to mechanosensory stimulation of the antennae [69]; [70,71]. This suggests that although the AHN pairs do not directly converge upon common downstream targets within the brain, the consequence of their coactivation would affect networks of neurons that converge further downstream upon the Giant Fiber circuit.

By spanning the brain and VNC, the AHNs innervate CNS structures that fundamentally differ in their functional architecture. Thus, the relative demographics targeted by AHN compartments may also differ between the brain and VNC. Similar to the brain, both AHN pairs predominantly target interneurons within the VNC, yet there were more glaring differences in the remaining neurons targeted by each pair (Figure 6H). A greater proportion of MsAHN output was directed towards motor neurons relative to the MtAHNs, whereas the MtAHNs had relatively greater downstream connectivity with descending and sensory neurons (MANC; Figure 6H, FANC; Figure S5D). There was virtually no convergence in the downstream targets between the AHN pairs (MANC; Figure 6I, FANC; Figure S5E) and while the output of individual MsAHNs converged upon a large number of targets, the output of the individual MtAHNs was more highly segregated. Finally, there was only moderate overlap in the neuropil occupied by either the presynaptic or postsynaptic sites of the MsAHN and MtAHN downstream partners that were within the top quartile of synapses per neuron group; the MsAHN partners largely occupied the wing and haltere neuropils, while the MtAHN partners largely occupied the leg neuropils (Figure S5F, G). This implies that within the VNC, not only do the AHN pairs mostly target separate populations of neurons, but that they also target different cell types within each network, suggesting fundamentally different functional organization.

To determine if, like in the brain, the populations of neurons targeted by each AHN pair converge upon common 2^nd^ order targets, we calculated the effective connection strength [72] between the AHNs and either all VNC motor neurons or VNC sensory neurons. Effective connection strength is a metric that serves as a proxy for the extent to which the neurons downstream of an AHN pair synaptically interact with each neuron within a reference neurons class. For instance, effective connection strength for motor neurons take into account 1) the proportion of input synapses from individual AHN pairs onto each interneuron, efferent or ascending neuron that subsequently synapses upon a given motor neuron, and 2) the proportion of input synapses from each interneuron to a given motor neuron out of all synapses received by that motor neuron (Figure S5H). In contrast to the brain, the populations of interneurons targeted by each AHN pair do not converge upon the same downstream 2nd order motor neuron targets (Figure 6J, Figure S5I). While the interneurons targeted by the MsAHNs predominantly synapse upon wing, haltere and neck motor neurons, the MtAHNs’ downstream interneuron population predominantly targets putative leg and abdominal motor neurons (Figure 6J, Figure S5I). To a lesser degree, this was also the case for the interneurons targeted by the AHNs that were upstream of sensory neurons (Figure 6K, Figure S5J). For instance, the MtAHNs targeted interneurons upstream of leg sensory neurons, while the MsAHNs targeted interneurons upstream of neck and haltere sensory neurons. However, unlike the effective connection strength for motor neurons, both pairs had some convergence upon interneurons that synapse upon wing sensory neurons (Figure 6K, Figure S5J). Thus overall, the 1^st^ and 2^nd^ order downstream targets of the AHN pairs remain largely segregated within the VNC, implying that each AHN pair is involved in distinct pre-motor circuits.

Having established that the two AHN pairs integrate considerable common input, but target separate downstream networks, we next sought to characterize the structure of these networks. We began by identifying the individual synaptic partners that receive the largest number of synapses from each AHN pair within MANC (Figure 7A, Figure S6A, B), although the MsAHNs in general made a larger number of synaptic contacts with these individual downstream partners relative to the MtAHNs (Figure 7). For the MsAHN, two groups of neurons received the greatest synaptic input; a population of wing-tectular interneurons (Figure 7B) and indirect flight wing motor neurons which innervate the dorsal longitudinal wing depressor muscles as well as the dorsal ventral muscles (Figure 7C). Interestingly, the wing-tectular interneurons and wing motor neurons formed a compact network that includes the DNg02s, DNp54s, and MsAHNs (Figure 7D). In this circuit, the cholinergic DNg02s likely provide fast excitatory input to AHNs via the nAChRα6 receptor expressed in both AHN types (Figure 5A). In addition to providing common input to all AHNs, the DNg02s and DNp54s synapse upon each other. The DNg02s also synapse heavily upon the wing tectular interneurons and the same wing motor neurons targeted by the MsAHNs. Finally, the wing-tectular interneurons heavily synapse upon the wing motor neurons targeted by the MsAHNs, creating a feedforward network originating with descending neurons known to regulate wing motor control [38,40]. In contrast to the MsAHNs, the downstream partners of the MtAHNs had very little interconnectivity (Figure S6C, D) and collectively consisted of a diverse set of ascending neurons, descending neurons and interneurons (Figure S6E-I).

**Figure 7.**
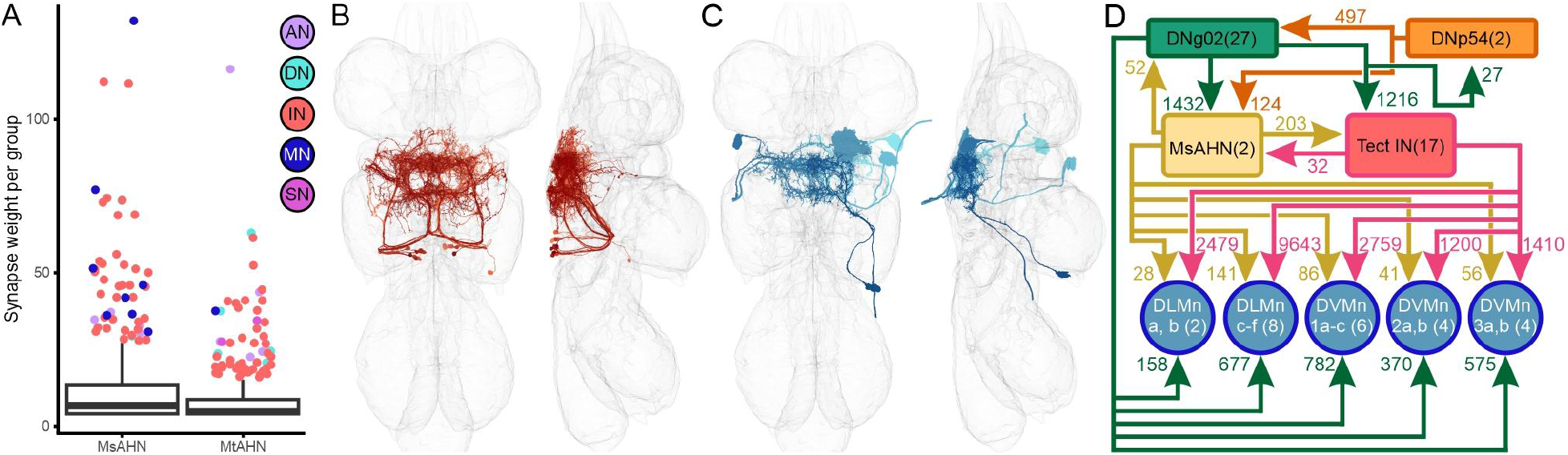
MsAHNs form a feedforward with the DNg02s to directly and indirectly target wing motor neurons. **A)** Box and whisker plot of AHN downstream partners for number of synapses per synaptic partner group (neurons with matching morphology) with the MsAHNs and MtAHNs. Outliers above the 3rd quartile plus 1.5 x interquartile range are plotted individually and color-coded based on cell class. **B-C)** The cell classes receiving the greatest amount of synaptic input from the MsAHNs were **B)** a population of tectular interneurons (shades of red) and **C)** wing motor neurons (shades of blue); only a subset of tectular interneurons and one of each wing motor neuron type are shown for clarity. **D)** Circuit motif depicting the relationship between the DNg02s (green), DNp54s (orange), MsAHNs (yellow), tectular interneurons (red) and wing MNs (blue). The DNg02s and DNp54s are reciprocally connected and both synapse upon the MsAHNs. The DNg02s and MsAHNs synapse upon the tectular interneurons and wing MNs (MNs of the dorsal longitudinal muscles; DLMn a, b and DLMn c-f, and of the dorsal ventral muscles; DVMn 1a-c, DVMn 2a, b and DVMn 3a, b).

## Discussion

Coordination between motor output and sensory processing is an essential feature of locomotion, at times requiring precise interactions between neuronal classes that each serve different roles and perform different computations. In this study, we comprehensively explored the synapse-level network architecture of histaminergic neurons that project broadly throughout the VNC and the brain. The AHNs appear to be a common feature of insect nervous systems [19], and the putative homologs of the MsAHNs in *Manduca sexta* provide a corollary discharge to the olfactory system, a function likely not conserved in *Drosophila* [4]. Furthermore, ascending CD neurons appear to be a common feature across the animal kingdom [2,73–75], therefore this study provides the opportunity to take a holistic perspective of how CDCs integrate into multiple networks to coordinate sensory and motor function.

Here, we reconstructed the AHNs and their upstream and downstream networks in EM volumes, revealing the comprehensive connectivity of the AHNs at the synapse level. However is the connectivity of the AHNs consistent with a CDC role or the closely-related concept of efference copy? By past definitions (reviewed in [76]; [2]; [77], corollary discharge generally refers to motor signals relayed to higher order motor or sensory circuits to modulate sensorimotor processing, planning or learning, while efference copy specifically refers to motor signals relayed to the early sensory system, often used to subtract reafferent sensory input. Circuit connectivity suggests that the MsAHNs relay a wing motor signal to visual motion processing circuits in the brain in a corollary discharge role, while they function in the VNC as a feedforward element in a flight circuit. In comparison, the MtAHNs integrate descending wing motor commands and relay them to putative leg networks within the VNC fitting a corollary discharge function, and to auditory efferents and first order auditory interneurons in the brain fitting an efference copy function.

In contributing to CDC-related roles, the two AHN pairs likely represent information about ongoing wing movement as they receive shared excitatory input from descending neurons involved in flight control, and we demonstrate that AHNs become active during flight, similar to *Manduca [4]*. Past work has demonstrated that the flight motor state is conveyed to several visual and visuomotor circuits in the brain [78–82], likely by multiple parallel pathways; this study is the first to document ascending pathways that convey flight motor information from the VNC to the brain. Indeed, the largest proportion of DN inputs to the AHNs derive from the DNg02s, which asymmetrically modulate wingbeat amplitude in response to widefield motion during flight [38,40]. In addition to the DNg02s, the DNp54s also provide shared input to both AHN pairs, and other DNs with as-yet-unknown functions provide distinct inputs to each AHN pair, suggesting that each AHN pair integrates distinct sets of other motor-related signals. These other motor signals are likely related to neck, wing and haltere motor control, or wing-leg coordination, as the axonal processes of these DNs largely target the upper, intermediate and lower tectulum of the VNC, which are associated with control of such behaviors [36–38]. Furthermore, upper tectulum DNs tend to receive input in the posterior slope of the brain, a neuropil implicated in visual processing and navigation, while intermediate and lower tectulum DNs are more diversely associated with several brain visuomotor processing centers and the antennal mechanosensory and motor center [36], suggesting that AHNs convey information related to visual and mechanosensory motor responses. While other neuron classes also provide input to AHNs, mapping of synapse locations of upstream cell classes onto AHN skeletons show that most DNs primarily synapse onto dendrites of the AHNs in the medial tectular regions of the nerve cord, while the other neuron classes largely synapse onto AHN axonal regions. The spatial distribution of synaptic input is thought to play roles in integration and computational processing of inputs [83], with axo-axonal synaptic input further playing roles in localized control of electrical transmission or neurotransmitter release [84]. Synapses from DNs onto the relatively simplistic AHN dendritic fields are likely closer to the spike initiation zone, suggesting that motor-related signals from DNs are the primary driver of AHN activity, consistent with both corollary discharge and efference copy functions, while other predominantly axo-axonal input may serve to locally refine AHN output, perhaps providing contextual information about the activity of downstream circuits to the local AHN axonal compartment.

In contrast to their shared upstream connectivity, each AHN pair appears to be functionally specialized in targeting distinct downstream networks, as their axonal processes tile largely non-overlapping regions in the VNC and brain, and each AHN pair contributes input to almost completely non-overlapping sets of neurons at the synapse level. In both the VNC and brain, interneurons comprise the bulk of downstream targets for both AHN pairs, while the remaining downstream connections (20-40%) show notable differences in cell class distribution. In particular, the MsAHNs more frequently synapse onto motor neurons in the VNC, while the MtAHNs more frequently synapse onto sensory afferents in the brain and VNC. This implies that the MtAHNs may play a role in impacting sensory detection, whereas the MsAHNs may in part impact motor output. From a network perspective, the MsAHNs are integrated into local flight circuits downstream of DNg02 and DNp54 and upstream of indirect flight motor neurons in the VNC, pointing towards a non-CDC feedforward role in VNC wing circuits. In the brain, MsAHNs synapse onto circuits that may play a role, at least in part, in wide-field visual motion processing and visuomotor responses [46–48], from their association with lobula plate tangential cells; this connectivity from motor output to sensorimotor processing centers is consistent with corollary discharge. As the DNg02s themselves activate during wide-field visual motion [38], it is possible that MsAHN downstream brain circuits further feed into DNg02s creating a motor output to sensorimotor processing feedback loop, although further connectomics work is required to examine this idea. Alternatively, MsAHN brain input may help suppress or modulate parallel visuomotor circuits while the fly is engaging in a flight-related motion response–MsAHN brain circuits do indeed have connectivity with DNa02, a major walking control DN [54].

In comparison, MtAHN downstream circuits in the VNC have much less overlap with their upstream circuits and are largely located in the leg neuropils, suggesting a role in modulating VNC leg motor output, although more work is required to dissect the circuit details and function of these leg neuropil circuits in general. In the brain, the MtAHNs target primary sensory afferents, notably the auditory JO-A and -B afferents of the Johnston’s organ (∼10% of synapses). Interestingly, at least half of all other MtAHN synapses in the brain are directed to neurons upstream or downstream of auditory afferents, overall suggesting that MtAHNs play a role in tuning or filtration of auditory signals, perhaps from self-generated acoustic signals related to wing movements. Many auditory neurons downstream of MtAHNs further provide input to the Giant Fiber descending neuron which drive a takeoff escape response [49–53], thus linking AHNs to at least one sound-responsive behavior. This arrangement has parallels to the sound-suppressing CDC of the cricket *Gryllus bimaculatus*, where a single pair of ascending neurons relays singing motor information from the VNC mesothoracic segment to auditory circuits in the prothoracic segment to suppress the auditory responses of sensory afferents and second order auditory neurons to self-generated courtship song [7,85,86]. While the arrangement of auditory organs fundamentally differs between flies and crickets, the common need for addressing auditory reafference has resulted in the evolution of at least superficially similar CDCs. It remains to be seen what similarities exist in the fly and cricket CDCs at synapse-level detail, and whether they have a common evolutionary origin.

In summary, we identified two pairs of corollary discharge neurons that span the brain and ventral nerve cord of *Drosophila* and provide insights into their circuit organization as well as their possible contributions to multiple sensorimotor functions. This lays the groundwork for future behavioral analyses of how AHNs contribute to shaping and tuning these sensorimotor functions including acoustic and visuomotor responses. From a wider perspective, the AHNs are but two pairs among roughly 1860 ANs (as surveyed in MANC), which likely function in relaying a wealth of information back to the brain including sensory, motor/behavioral, and other internal states. Significant headway has been made in understanding a handful of ANs, which play a range of roles from sensory detection to adaptive motor control [87–96]. Crucially, a broad behavioral survey and analysis of the activity of ∼250 AN types during behavior suggest that most function in encoding and relaying information to the brain about high-level behavioral states such as walking and grooming, rather than low-level states such as individual limb movement [96]. These functions may be carried out by integration and processing of multiple sources of proprioceptive or other sensory input, or by corollary discharge from VNC motor circuits as demonstrated by the AHNs. If the results of this survey are representative of AN function at large in *Drosophila*, corollary discharge functions may be widespread among ANs and constitute an open field for further study.

## Materials and Methods

### Fly stocks

All fly stocks were raised on a standard cornmeal/agar/yeast medium at 24°C on a 12:12 light/dark cycle at ∼60% humidity. For optogenetic stimulation of DNg02 by CsChrimson with 2-photon calcium imaging of MtAHNs, parental flies were allowed to lay eggs on standard cornmeal media containing 0.2 mM all-trans retinal. Newly-eclosed offspring (≤1 day post-eclosion) were transferred to cornmeal media containing 0.4 mM all-trans retinal and aged for 3-4 days before use in experiments. All fly cultures containing all-trans retinal were shielded from light. The fly stocks used in this study are summarized in Tables 1 and 2.

**Table 1:**
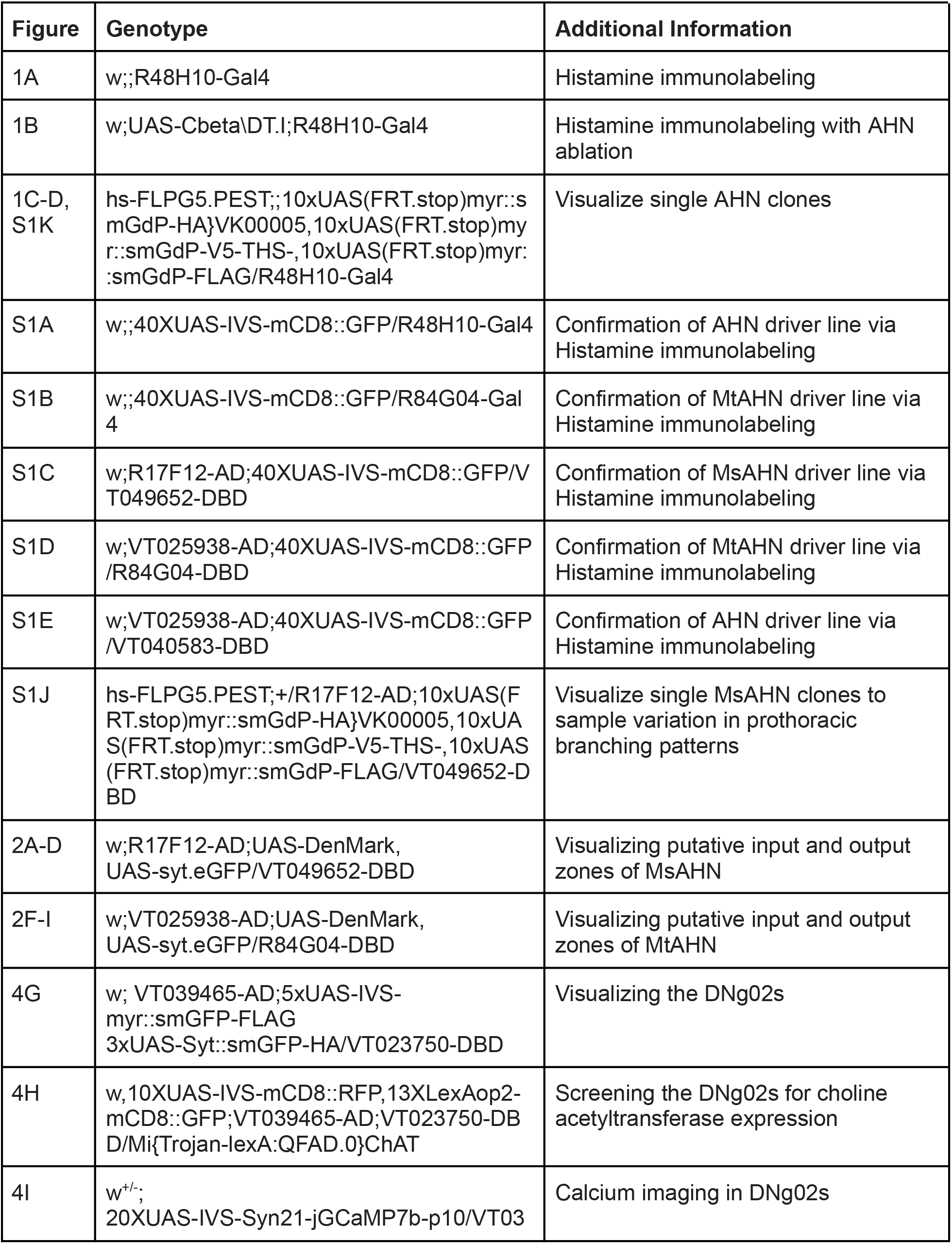
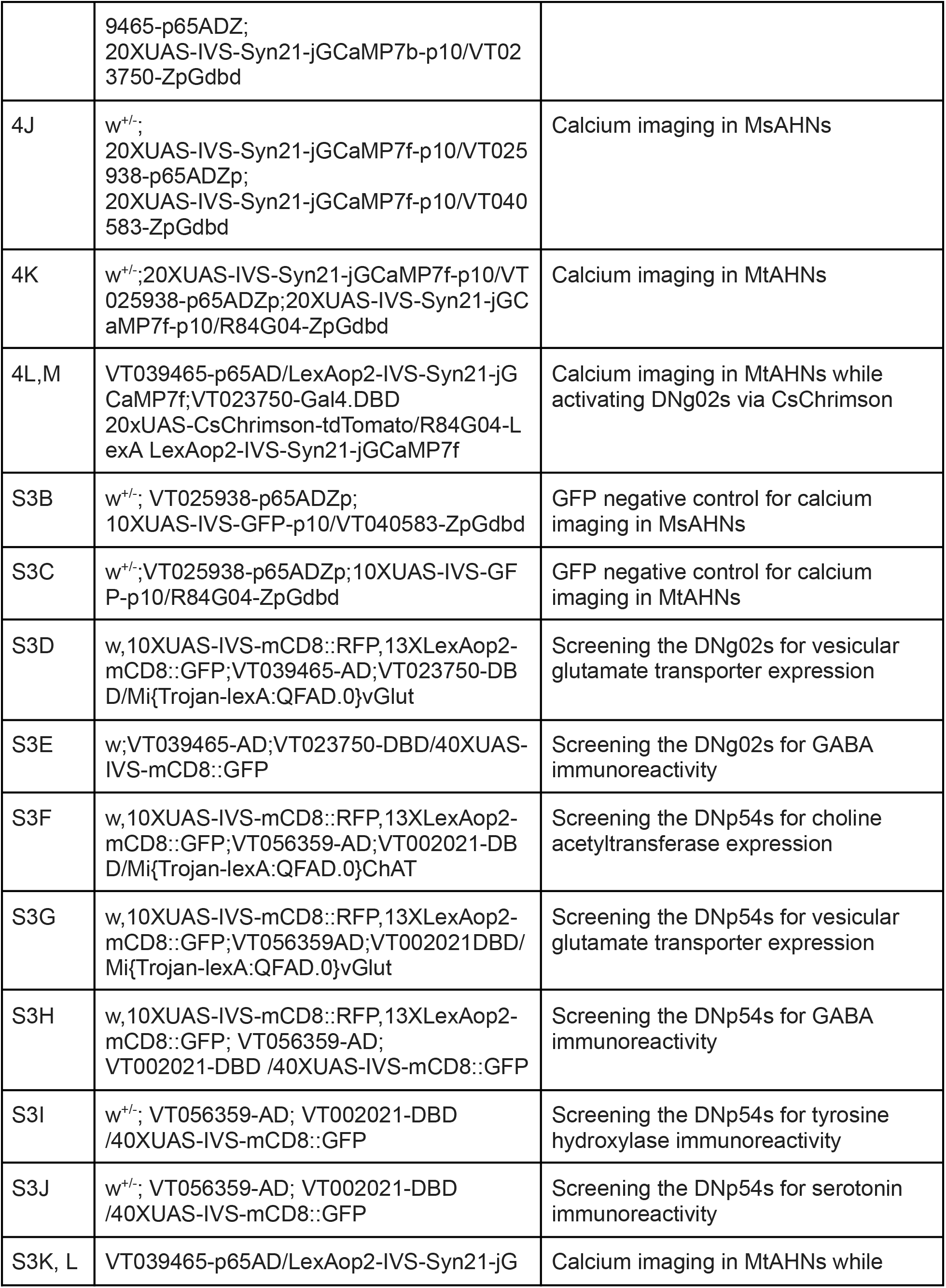
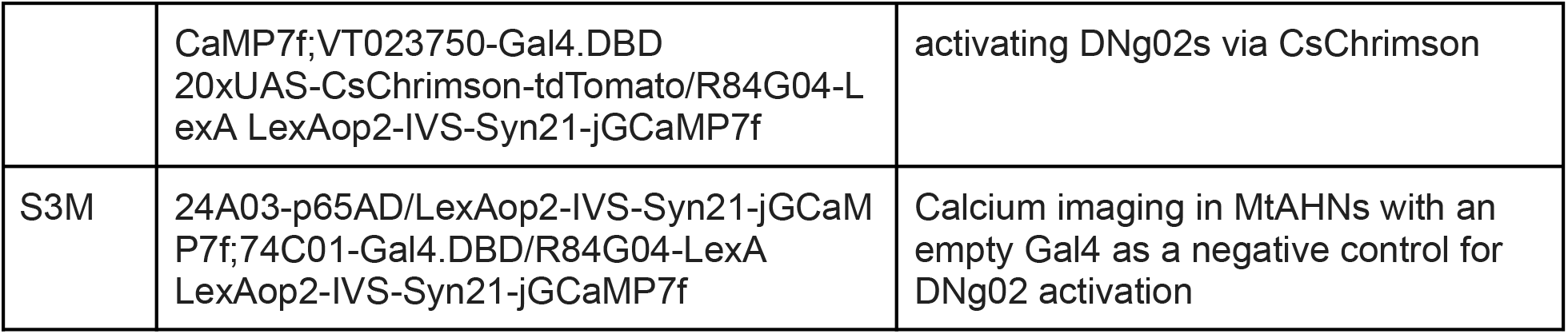
Genotype of flies in each figure

| Figure | Genotype | Additional Information |
| --- | --- | --- |
| 1A | w;;R48H10-Gal4 | Histamine immunolabeling |
| 1B | w;UAS-Cbeta<sup>DT.I</sup>;R48H10-Gal4 | Histamine immunolabeling with AHN ablation |
| 1C-D, S1K | hs-FLPG5.PEST;;10xUAS(FRT.stop)myr::smGdP-HA}VK00005,10xUAS(FRT.stop)myr::smGdP-V5-THS-,10xUAS(FRT.stop)myr::smGdP-FLAG/R48H10-Gal4 | Visualize single AHN clones |
| S1A | w;;40XUAS-IVS-mCD8::GFP/R48H10-Gal4 | Confirmation of AHN driver line via Histamine immunolabeling |
| S1B | w;;40XUAS-IVS-mCD8::GFP/R84G04-Gal4 | Confirmation of MtAHN driver line via Histamine immunolabeling |
| S1C | w;R17F12-AD;40XUAS-IVS-mCD8::GFP/VT049652-DBD | Confirmation of MsAHN driver line via Histamine immunolabeling |
| S1D | w;VT025938-AD;40XUAS-IVS-mCD8::GFP/R84G04-DBD | Confirmation of MtAHN driver line via Histamine immunolabeling |
| S1E | w;VT025938-AD;40XUAS-IVS-mCD8::GFP/VT040583-DBD | Confirmation of AHN driver line via Histamine immunolabeling |
| S1J | hs-FLPG5.PEST;+/R17F12-AD;10xUAS(FRT.stop)myr::smGdP-HA}VK00005,10xUAS(FRT.stop)myr::smGdP-V5-THS-,10xUAS(FRT.stop)myr::smGdP-FLAG/VT049652-DBD | Visualize single MsAHN clones to sample variation in prothoracic branching patterns |
| 2A-D | w;R17F12-AD;UAS-DenMark, UAS-syt.eGFP/VT049652-DBD | Visualizing putative input and output zones of MsAHN |
| 2F-I | w;VT025938-AD;UAS-DenMark, UAS-syt.eGFP/R84G04-DBD | Visualizing putative input and output zones of MtAHN |
| 4G | w; VT039465-AD;5xUAS-IVS-myr::smGFP-FLAG<br>3xUAS-Syt::smGFP-HA/VT023750-DBD | Visualizing the DN<sub>g</sub>02s |
| 4H | w,10XUAS-IVS-mCD8::RFP,13XLexAop2-mCD8::GFP;VT039465-AD;VT023750-DBD/Mi{Trojan-lexA:QFAD.0}ChAT | Screening the DN<sub>g</sub>02s for choline acetyltransferase expression |
| 4I | w<sup>+/-</sup>;<br>20XUAS-IVS-Syn21-jGCaMP7b-p10/VT03 | Calcium imaging in DN<sub>g</sub>02s |
|  | 9465-p65ADZ;<br>20XUAS-IVS-Syn21-jGCaMP7b-p10/VT023750-ZpGdbd |  |
| 4J | w<sup>+/-</sup>;<br>20XUAS-IVS-Syn21-jGCaMP7f-p10/VT025938-p65ADZp;<br>20XUAS-IVS-Syn21-jGCaMP7f-p10/VT040583-ZpGdbd | Calcium imaging in MsAHNs |
| 4K | w<sup>+/-</sup>;20XUAS-IVS-Syn21-jGCaMP7f-p10/VT025938-p65ADZp;20XUAS-IVS-Syn21-jGCaMP7f-p10/R84G04-ZpGdbd | Calcium imaging in MtAHNs |
| 4L,M | VT039465-p65AD/LexAop2-IVS-Syn21-jGCaMP7f;VT023750-Gal4.DBD<br>20xUAS-CsChrimson-tdTomato/R84G04-LexA LexAop2-IVS-Syn21-jGCaMP7f | Calcium imaging in MtAHNs while activating DNg02s via CsChrimson |
| S3B | w<sup>+/-</sup>; VT025938-p65ADZp;<br>10XUAS-IVS-GFP-p10/VT040583-ZpGdbd | GFP negative control for calcium imaging in MsAHNs |
| S3C | w<sup>+/-</sup>;VT025938-p65ADZp;10XUAS-IVS-GFP-p10/R84G04-ZpGdbd | GFP negative control for calcium imaging in MtAHNs |
| S3D | w,10XUAS-IVS-mCD8::RFP,13XLexAop2-mCD8::GFP;VT039465-AD;VT023750-DBD/Mi{Trojan-lexA:QFAD.0}vGlut | Screening the DNg02s for vesicular glutamate transporter expression |
| S3E | w;VT039465-AD;VT023750-DBD/40XUAS-IVS-mCD8::GFP | Screening the DNg02s for GABA immunoreactivity |
| S3F | w,10XUAS-IVS-mCD8::RFP,13XLexAop2-mCD8::GFP;VT056359-AD;VT002021-DBD/Mi{Trojan-lexA:QFAD.0}ChAT | Screening the DNp54s for choline acetyltransferase expression |
| S3G | w,10XUAS-IVS-mCD8::RFP,13XLexAop2-mCD8::GFP;VT056359AD;VT002021DBD/Mi{Trojan-lexA:QFAD.0}vGlut | Screening the DNp54s for vesicular glutamate transporter expression |
| S3H | w,10XUAS-IVS-mCD8::RFP,13XLexAop2-mCD8::GFP; VT056359-AD; VT002021-DBD /40XUAS-IVS-mCD8::GFP | Screening the DNp54s for GABA immunoreactivity |
| S3I | w<sup>+/-</sup>; VT056359-AD; VT002021-DBD /40XUAS-IVS-mCD8::GFP | Screening the DNp54s for tyrosine hydroxylase immunoreactivity |
| S3J | w<sup>+/-</sup>; VT056359-AD; VT002021-DBD /40XUAS-IVS-mCD8::GFP | Screening the DNp54s for serotonin immunoreactivity |
| S3K, L | VT039465-p65AD/LexAop2-IVS-Syn21-jG | Calcium imaging in MtAHNs while |
|  | CaMP7f;VT023750-Gal4.DBD<br>20xUAS-CsChrimson-tdTomato/R84G04-LexA LexAop2-IVS-Syn21-jGCaMP7f | activating DNg02s via CsChrimson |
| S3M | 24A03-p65AD/LexAop2-IVS-Syn21-jGCaMP7f;74C01-Gal4.DBD/R84G04-LexA LexAop2-IVS-Syn21-jGCaMP7f | Calcium imaging in MtAHNs with an empty Gal4 as a negative control for DNg02 activation |

**Table 2:**
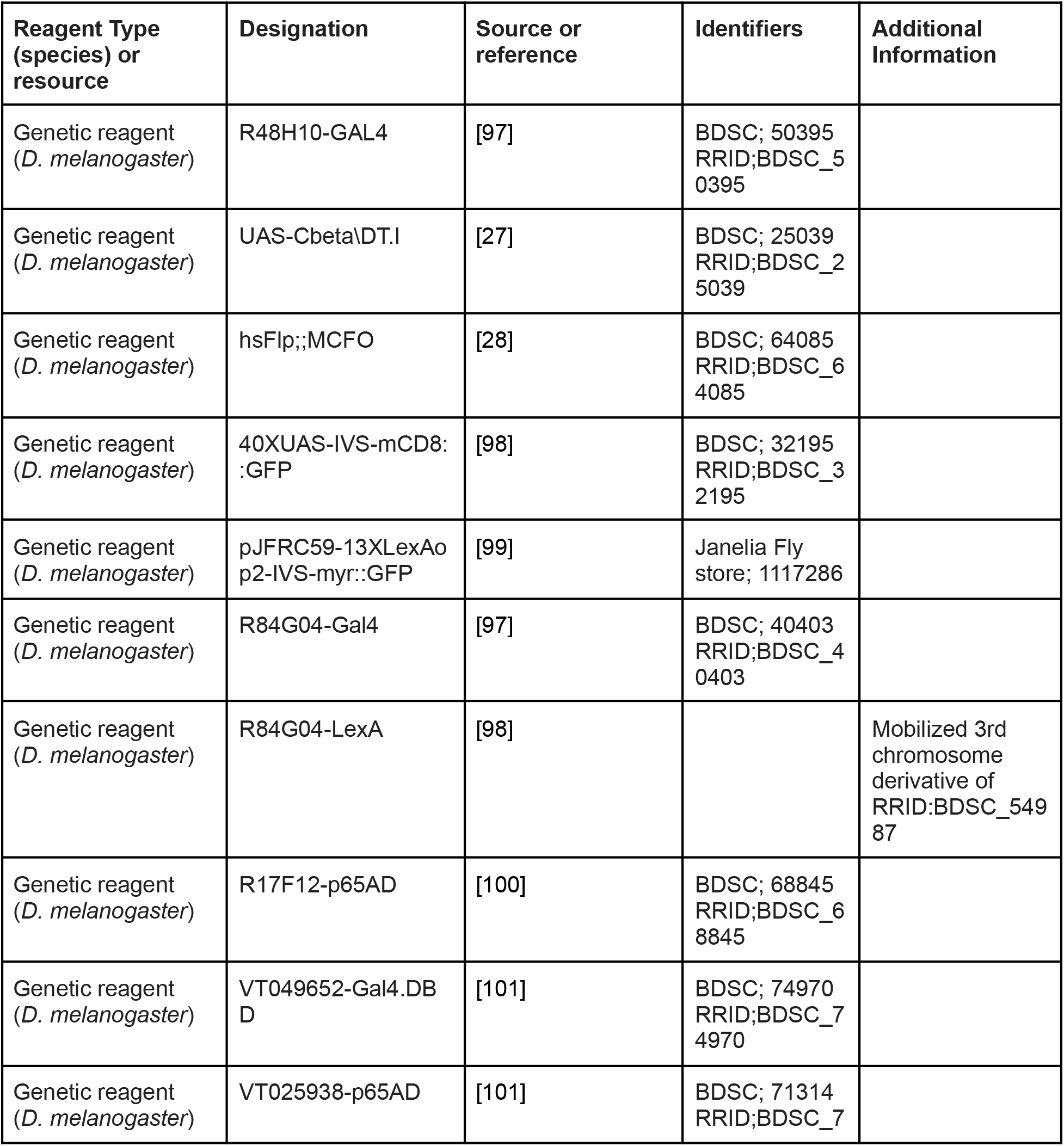
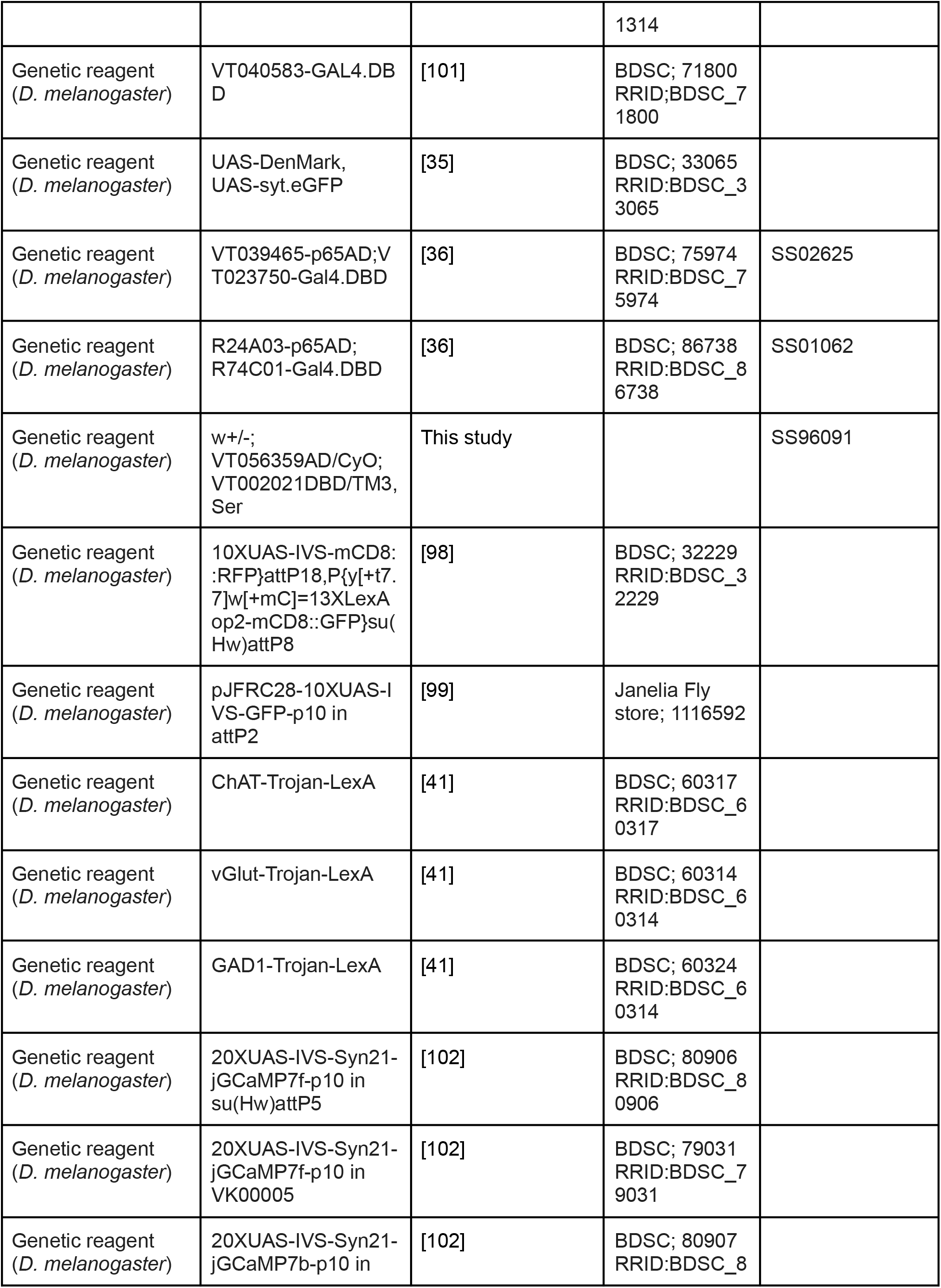
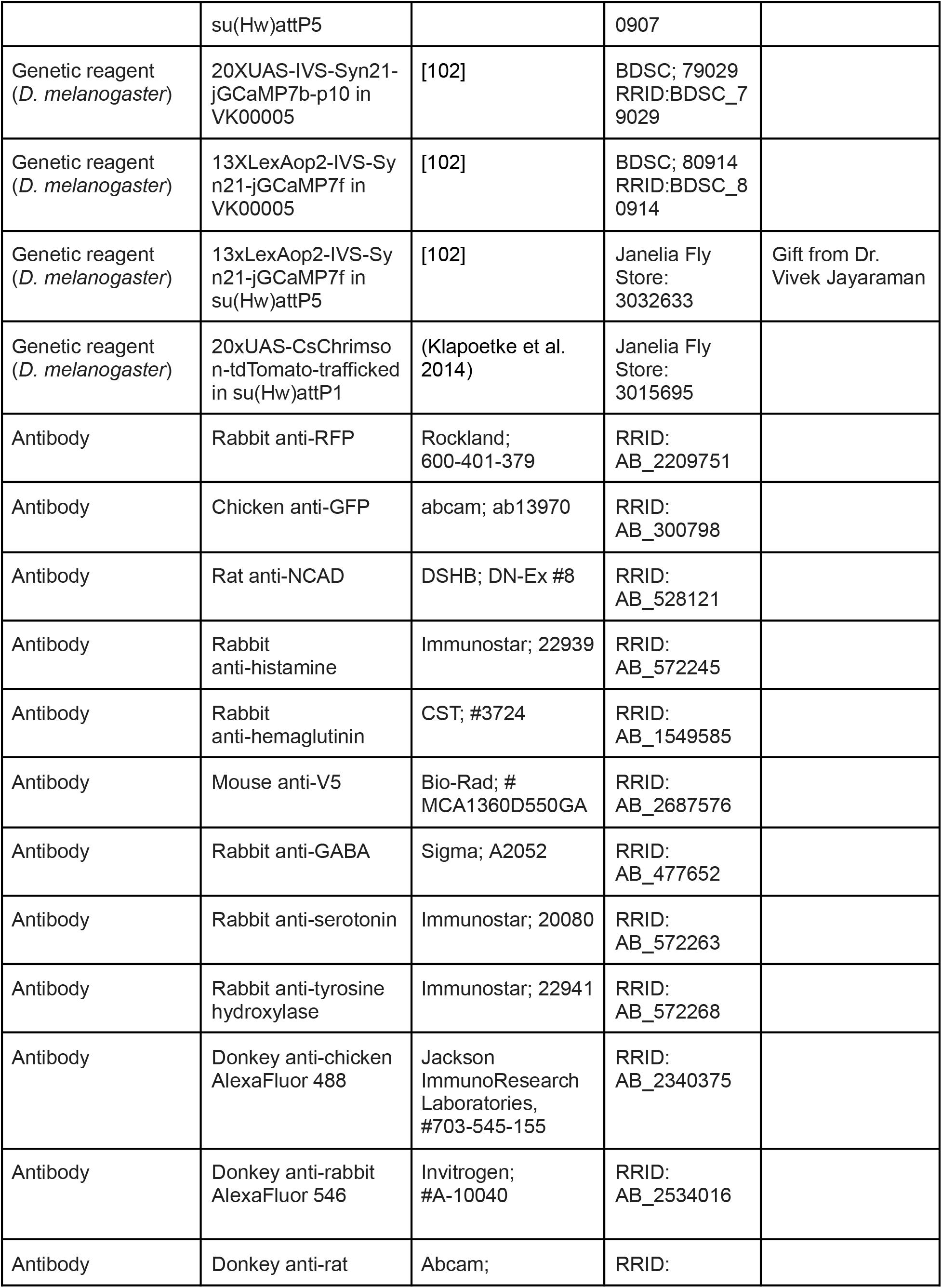
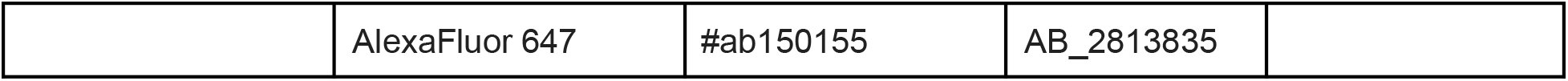
Key Resources and Reagents

### Immunohistochemistry

Intact brain and ventral nerve cords were dissected in *Drosophila* saline [103] and fixed in 4% paraformaldehyde (PFA) at 4°C for 30 minutes, unless immunostaining for histamine in which samples were fixed in 4% 1-Ethyl-3-(3-Dimethylaminopropyl)carbodiimide in PBS at 4°C for 2 hours before post-fixing in 4% PFA at 4°C for 30 minutes. Samples were then washed 4X in PBST (PBS with 0.5% Triton X-100) and blocked for 1 hour in 2% BSA (in PBST and 50mM sodium azide), except when labeling for histamine in which 3% normal goat serum (in PBST and 50mM sodium azide) was used as the blocking agent. Primary antibodies (see Table 2 for antibody details) were applied for 48 hours at 4°C with agitation. After, samples were washed 4X in PBST and blocked as described above. Secondary antibodies were applied and incubated for 48 hours in 4°C with agitation. Samples were then washed 2X in PBST and 2X in PBS before being run through an ascending glycerol series (40%, 60% and 80%) for 10 minutes each. Samples were mounted in VectaShield. Images were analyzed with an Olympus FV1000 BX61 (Shinjuku, Tokyo, Japan) confocal, using Fluoview FV1000 software with a 20X UPlanSApo, 40x UPlanFL-N or 60x PlanApo-N oil-immersion objective.

### Transcriptomic analysis

Initial exploration of the adult single-cell VNC transcriptomic atlas [43] was done using the online SCope tool (scope.aertslab.org). To narrow down candidate ascending histaminergic neurons (AHNs), we used the the SCope online tool to select cells that express the histidine decarboxylase (*Hdc*) gene within the aminergic cell cluster identified in [43], and downloaded the gene expression dataset for those cells. Next, we examined expression of the hox genes antennapedia (*Antp*) and ultrabithorax (*Ubx*) to locate cells likely to be found in the mesothoracic and metathoracic neuromeres, respectively. Using an R script with the packages loomR (https://github.com/mojaveazure/loomR), gplots (https://github.com/talgalili/gplots) and viridis (https://github.com/sjmgarnier/viridis), we selected a subset of cells expressing *Hdc* and one of *Antp* and *Ubx* with raw read counts of ≥2, and further narrowed down cells where the raw expression of one of Antp and Ubx is much greater than the other (in practice, ≥15:1). Then, we normalized all gene expression levels by counts per million (CPM) followed by log2(n+1) scaling. The likely AHNs, their cell IDs within the transcriptome dataset, and their normalized expression of *Hdc*, *Antp* and *Ubx* are shown in Table 3. Neuronal genes of interest covering neurotransmitter receptors, second messenger proteins, SNARE proteins and ion channels were then selected and plotted by heatmap. For each gene set, we then generated gene expression heatmaps and hierarchical clustering of genes and cells using the complete linkage method on Euclidean distance.

**Table 3:**
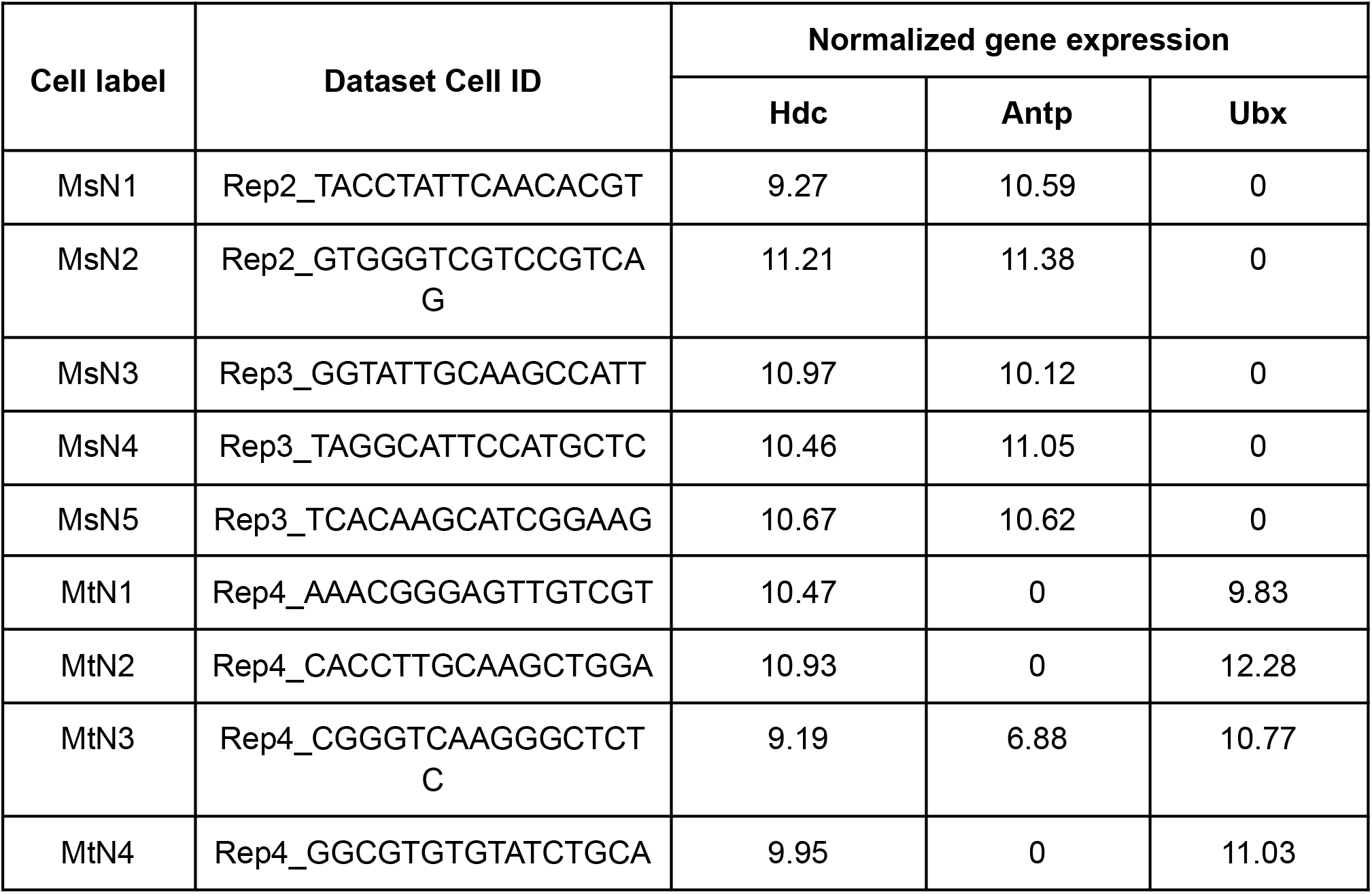
Cell IDs in Scope dataset used for transcriptomic analysis

### Circuit reconstruction and connectomic analyses

The AHNs were reconstructed and their pre- and postsynaptic sites annotated in the Female Adult Nerve Cord (“FANC”) dataset [30,31] and the Female Adult Fly Brain (“FAFB”) datasets [29] using CATMAID (Collaborative Annotation Toolkit for Massive Amounts of Image Data) [104,105]. We further identified the AHNs in the Male Adult Nerve Cord (“MANC”) by their connectivity [32,33]. We identified the following cells as the top AHN candidates in VNC and brain EM volumes: For MsAHNs, skeleton IDs 237078 and 402598 in FANC CATMAID (segment IDs 648518346499994886 and 648518346489573207 in FANC autosegmentation), bodyIDs 13926 and 12536 in MANC, and skeleton IDs 2455455 and 2455571 in FAFB CATMAID; for MtAHNs, skeleton IDs 313368 and 250373 in FANC CATMAID (segment IDs 648518346488561230 and 648518346475813602 in FANC autosegmentation), bodyIDs 42819 and 11003 in MANC, and skeleton IDs 3385431 and 17138817 in FAFB CATMAID. Pre- and postsynaptic partners were reconstructed either manually or using autosegmentation AI developed for FAFB [106] and FANC [30]. For FANC, we reconstructed all presynaptic partners of the AHNs, while postsynaptic partners were reconstructed in the autosegmentation volume only for bodies with an initial synapse count of ≥3 with any AHN. For FAFB, we reconstructed postsynaptic partners for ∼50% of downstream connections of MsAHN-L and MtAHN-L, not counting orphan fragments. For MANC, we retrieved upstream and downstream connectivity of AHNs from the MANC neuprint server (https://neuprint.janelia.org/?dataset=manc) and further filtered connectivity only with ‘valid’ neurons (has a neuron class entry in the neuprint ‘class’ field). Traced or annotated upstream and downstream connection counts of AHNs in all 3 EM volumes are summarized in Table 4. The automated synapse predictions in MANC yielded larger upstream and downstream synapse counts for AHNs compared to manual synapse annotations in FANC, possibly due to differences in EM volume preparation and imaging, most notably lower T-bar staining intensity in FANC as well as higher voxel resolution in the Z-plane for MANC (8×8×8 nm^3^) compared to FANC (4.3×4.3×45 nm^3^).

**Table 4:** FANC, MANC and FAFB AHN tracing and connectivity summary

| **EM volume** | **Cell** | **Postsynaptic sites/upstream connections (to valid neurons)** | **Presynaptic sites** | **Downstream connections (to valid neurons/orphans/undetermined)** |
| --- | --- | --- | --- | --- |
| FANC | MsAHN-L | 1040 (992) | 1664 | 4604 (995/93/3516) |
|  | MsAHN-R | 962 (899) | 1170 | 3183 (740/60/2383) |
|  | MtAHN-L | 1430 (1288) | 2586 | 7580 (1317/505/5758) |
|  | MtAHN-R | 1024 (935) | 1952 | 4901 (891/292/3718) |
| MANC | MsAHN-L | 2941 (2825) | 1782 | 5090 |
|  | MsAHN-R | 3116 (2984) | 1722 | 5001 |
|  | MtAHN-L | 4039 (3499) | 2102 | 4946 |
|  | MtAHN-R | 4895 (4173) | 2549 | 6032 |
| FAFB | MsAHN-L | 92 | 1115 | 3427 (1649/407/1371) |
|  | MtAHN-L | 70 | 556 | 1440 (812/215/413) |

Data analysis and visualization was carried out with R version 4.1.1 with the following packages: fancr (https://github.com/flyconnectome/fancr), fafbseg (https://github.com/natverse/fafbseg), reticulate (https://github.com/rstudio/reticulate), googlesheets4 (https://github.com/tidyverse/googlesheets4), neuprintr (https://natverse.org/neuprintr/), malevnc (https://github.com/natverse/malevnc), nat [107] and nat-nblast [34], catmaid (https://github.com/natverse/rcatmaid), igraph (https://github.com/igraph/igraph), ggplot2 (https://github.com/tidyverse/ggplot2), and plyr (https://github.com/hadley/plyr).

We utilized NBLAST [34] in two ways, first as a method to determine if neurons with similar morphologies to the 2 pairs of AHNs were present in the FANC and MANC volumes, and second to match DN types connected to the AHNs in FANC to DNs in MANC. For finding cells with similar morphology to the AHNs in MANC, we retrieved skeletons for our top pair of MsAHN and MtAHN candidates, as well as skeletons for all MANC neurons with soma in T2 or T3. Skeletons were rescaled to micron dimensions, small neurites less than 10 μm long were pruned to reduce spurious short-length projections resulting from skeletonization methods, and skeletons were then registered to a symmetrized MANC template provided via the malevnc package. We carried out NBLAST via the R nat-nblast package [34] using the version 2 algorithm optimized for *Drosophila* neurons, for each of the two MsAHN and MtAHN candidates against the set of MANC neurons with soma in T2 or T3, respectively. To determine the top 5 matches against the MsAHN and MtAHN cell types, we pooled the top hits for each pair of cells, removed redundant matches between the cell pairs and self-matches, then took the top 5 cells rank-ordered by NBLAST score.

To determine FANC DN types, we skeletonized neuron meshes (wavefront method) for DNs upstream and downstream of AHNs from the FANC autosegmentation volume via the skeletor python package (https://github.com/navis-org/skeletor). FANC skeletons were registered to MANC space via a FANC-to-MANC registration provided in the R malevnc package. For MANC DNs, we retrieved skeletons for all DNs from the MANC neuprint server (https://neuprint.janelia.org/). All skeletons were rescaled to micron dimensions, and small neurites less than 10 μm long were pruned. NBLAST between each FANC DN and the entire set of MANC DNs (plus the FANC DN itself for normalization of NBLAST scores) was carried out with the version 2 NBLAST algorithm. The top 10 hits from MANC were then manually compared with the reference neuron from FANC to verify matches.

We used CATMAID or custom R code to generate synapse fraction and connectivity graph plots. For FANC and FAFB, we used a synapse count of 3 as a minimum threshold for defining significant synaptic partners of AHNs. Upstream and downstream synapse counts are higher in MANC compared to FANC. Thus, to establish an equivalent threshold, we used the ratio of upstream connections provided by ‘valid’ neurons (not counting orphan fragments) for all AHNs in MANC to that of FANC, multiplied by the FANC synapse threshold and rounded, as the MANC upstream threshold. For the MANC downstream threshold, we used the sum of AHN downstream synapses in FANC provided by both valid neurons and undetermined bodies for calculating this ratio, as we had selectively reconstructed and proofread only bodies that had an initial downstream synapse count of 3 or more with AHNs. These calculations yielded an upstream synapse threshold of 10 and downstream synapse threshold of 3 for MANC AHNs. To generate the synapse distribution figures, we plotted the xy and xz coordinates of the locations of the skeleton nodes and of the input synapses of the left AHNs from FANC CATMAID.

To summarize AHN upstream connectivity with DNs, we categorized upstream DNs above the established synapse thresholds by their cell type, as determined by curation in MANC by the Jefferis group [39] and our NBLAST of FANC DNs to MANC DNs. FANC DN identities were further corroborated with the Jefferis group (personal communication). We defined the most significant DN inputs to the AHNs as contributing ≥5% of DN input by type-to-type connectivity (that is, by sum of connectivity for all DNs of the same type to all AHNs of the same type). DNs below this threshold were placed into the “DN (summed)” category for plotting of DN synapse fractions and graphs.

To calculate the effective connection strength [72] from the AHNs to VNC motor neurons or sensory neurons in MANC, we retrieved synapse connectivity between all valid neurons in MANC, collapsed connectivity by cell type (all neurons except sensory) or subclass (grouping by anatomical origin and cell morphology for sensory neurons), and converted synapse weights to input fractions. For each motor neuron type or sensory neuron subclass, we calculated effective connection strength for a path length of 2 (one synaptic hop) downstream of AHNs. That is, for each possible intermediary neuron, the input fraction contributed by AHNs to this neuron is multiplied by the input fraction contributed by the neuron to the motor neuron type or sensory neuron subclass; the effective connection strength of an AHN to the motor neuron type or sensory neuron subclass is the sum of these multiplied numbers for all intermediary neurons. DNs or sensory neurons were not considered for intermediary neurons, as they either receive input outside the VNC or are driven by non-neuronal inputs.

For ROI input and output of AHN downstream partners in MANC, we retrieved ROI presynaptic and postsynaptic site counts for these partners, combined counts for left and right neuropils and normalized them by their totals for each neuron. The per-neuropil mean for each neuron type was then calculated. For MNs, we set their presynaptic site count to zero, as manual examination in the EM volume suggested that most predicted sites for these MNs are false positives.

### Calcium Imaging and analysis

Imaging of flies was carried out on a custom-built two-photon/epifluorescence microscope. To image AHN somas during flight, we cold anesthetized flies, removed legs and mounted flies ventral side up on custom-made fly holders [108] with Loctite AA 3972 (Part# 36294) light-activated glue. We dissected through the ventral side of the thorax in external saline (103 mM NaCl, 3 mM KCl, 5 mM TES, 8 mM trehalose-2H_2_O, 10 mM glucose, 26 mM NaHCO_3_, 1 mM NaH_2_PO_4_, 1.5 mM CaCl_2_-2H_2_O, 4 mM MgCl_2_, 270-275 mOsm) [109] saturated with carbogen (95% O2/5% CO2). We used epifluorescence imaging to measure calcium activity of AHNs during flight, as thoracic vibrations caused by flight affected capture of the optically-sectioned 2-photon image stacks. Epifluorescence imaging of AHN somata was carried out through a Nikon CFI75 LWD 16X W objective with a pco.edge 5.5 monochrome camera (pco.) at 20 Hz using µManager software [110], with excitation by a 470 nm LED source, and a filter set consisting of a 495 nm dichroic beamsplitter (Semrock), 447/60 nm excitation (Semrock) and 525/80 nm emission filter (MidOpt). The behavior of the fly was simultaneously monitored with a Blackfly S BFS-U3-04S2M-CS monochrome camera (FLIR) with a 845/60 nm bandpass filter (MidOpt) under near-IR illumination. To induce flight, an airpuff was delivered to the head ∼10 seconds into the imaging period, and flies were imaged for an additional 20 seconds. Flight trials were carried out 3 times per fly. Six to nine flies which flew robustly for all 3 trials were used for further analysis.

For imaging of DNg02 dendrites during flight, we head-fixed flies and dissected through the posterior surface of the head. Two-photon imaging was carried out with a 920 nm Insight DS+ pulsed laser (MKS Instruments) at 7-8.5 mW power, with emission detected through a Nikon CFI75 LWD 16X W objective by a photomultiplier tube (Hamamatsu H10770PB-40-SEL) through a 503/40 nm dichroic filter (Semrock). Volume imaging was carried out at a rate of 14.14 Hz (10 z-slices with 5 μm increments). Microscope control and image capture was controlled through Scanimage software version 2016 [111]. Induction of flight and monitoring of fly behavior was carried out as for imaging of AHN somata. Flight trials were carried out 3 times per fly. Six flies which flew robustly for all 3 trials were used for further analysis.

For imaging of MtAHNs with optogenetic stimulation of DNg02s, flies were mounted ventral side up and dissected through the ventral thorax as for imaging of AHN soma. Two-photon imaging was carried out for MtAHN soma as for imaging of DNg02 dendrites above. To carry out optogenetic stimulation of DNg02s, red light from a 660 nm LED source filtered through a 661/20 nm dichroic filter (Semrock) was presented for one second through the objective. Optogenetic stimulation was carried out 3 times per fly. Six flies per experimental condition were tested.

Image analysis for AHNs and DNg02s was carried out in Fiji ver 2.3.0/1.53f51 [112]. Image stacks acquired using 2-photon microscopy were first flattened via sum slices projection. For AHN flight experiments, motion in the VNC was observed during flight, which we corrected using a rigid motion correction method implemented in the moco ImageJ plugin [113]. ROIs were then manually drawn around the AHN somas or DNg02 dendrites, as well as in an empty area of the neuropil around the soma for background subtraction, and the signal intensity within these ROIs were measured across time. Further data processing and figure generation was carried out in MATLAB ver 2019b. Baseline fluorescence was measured in the 5-second period before flight induction and used to normalize fluorescence intensity changes across each calcium trace. If more than one soma was observable per fly, quantification and plotting was carried out separately for each soma.

### Resource availability

#### Lead contact

Further information and requests for resources and reagents should be directed to and will be fulfilled by the lead contact, Andrew M. Dacks.

### Materials availability

SplitGal4, Gal4 and LexA *Drosophila* stocks generated in this paper are available from the lead contact without restriction upon request.

### Data and code availability

All data and code are available on request from the lead contact.

### Author contributions

Conceptualization, A.M.D., K.C.D. and G.M.C.; Methodology, H.S.J.C., K.N.B., K.C.D. and A.M.D.; Formal analysis, H.S.J.C. and K.N.B.; Investigation, H.S.J.C., K.N.B, M.M.D., F.S., J.D.R., K.H., R.F.A., A.M.P., A.P.C., M.E., K.C.D. and A.M.D.; Resources, J.S.P. and W.C.A.L.; Writing–Original Draft, H.S.J.C., K.N.B., K.C.D. and A.M.D.; Writing–Review & Editing, J.S.P., A.P.C., M.E. and W.C.A.L.; Visualization, H.S.J.C., K.N.B., F.S. and A.M.D.; Supervision, A.M.D., K.C.D. and G.M.C.; Funding Acquisition, A.M.D., K.C.D. and G.M.C.

**Figure S1.**
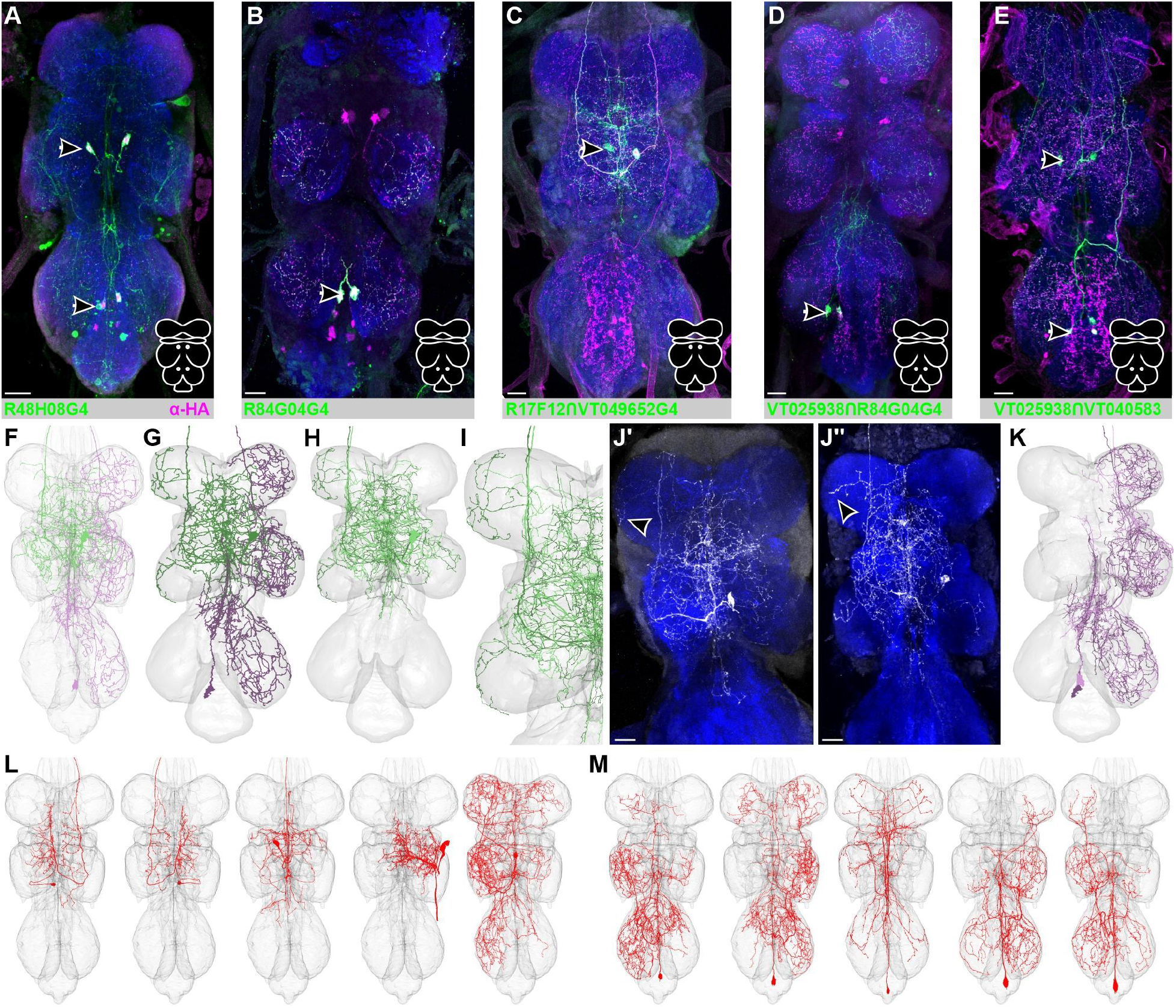
Validation of driver lines including the AHNs driving expression of GFP (green) and immunolabeled for histamine (magenta) NCAD is used as a neuropil marker (dark blue). Cartoon schematic at lower right corner indicates which AHN pairs are expressed by a given driver line. **A)** R48H10-Gal4 (both AHN pairs included). **B)** R84G04-Gal (MtAHNs included). **C)** R17F12 ∩ VT049652 splitGal4 (MsAHN only). **D)** VT025938 ∩ R84G04 splitGal4 (MtAHNs only). **E)** VT025938 ∩ VT049652 splitGal4 (both AHN pairs). **F-G)** reconstructions of the MsAHN (green) and MtAHN (magenta) in the **F)** MANC and **G)** FANC datasets. **H)** Overlaid reconstructions of a single MsAHN from the MANC (light green) and FANC (dark green) datasets. **I)** The processes of MsAHNs within the prothoracic neuromere differ between MANC and FANC in terms of the extent to which they project laterally. **J)** Single MCFO clones of a male MsAHN showing the variability in the prothoracic branching consistent with differences between FANC and MANC. **K)** Overlaid reconstructions of a single MtAHN from the MANC (light magenta) and FANC (dark magenta) datasets revealed no obvious morphological differences. **L)** Top 5 non-redundant NBLAST hits from all neurons with soma in T2 in the MANC dataset queried against the pair of MsAHN candidates. Body IDs from left to right are 11628, 12486, 10378, 10178 and 10222. **M)** Top 5 NBLAST hits from all neurons with soma in T3 in the MANC dataset queried against the pair of MtAHN candidates. Body IDs from left to right are 10197, 10330, 10066, 10104 and 10109. All scale bars = 20μm.

**Figure S2.**
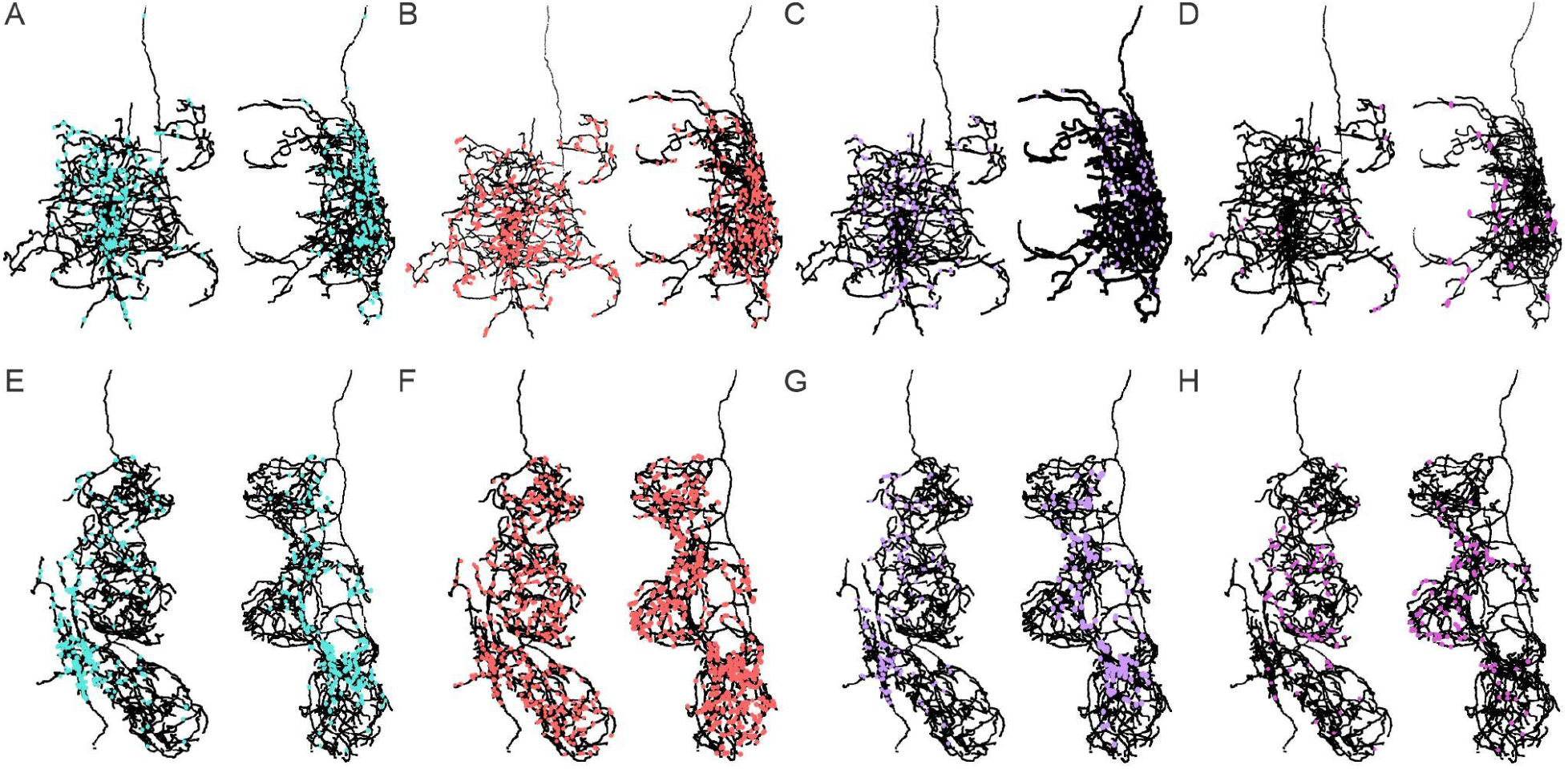
Input synapse distribution on AHNs by cell class. **A-D)** Horizontal (left) and sagittal (right) views of the synapse distributions upon the left MsAHN from **A)** descending neurons (cyan), **B)** interneurons (red), **C)** ascending neurons (lavender) and **D)** sensory neurons (pink). **E-H)** Horizontal (left) and sagittal (right) views of the synapse distributions upon the left MtAHN from **E)** descending neurons (cyan), **F)** interneurons (red), **G)** ascending neurons (lavender) and **H)** sensory neurons (pink).

**Figure S3.**
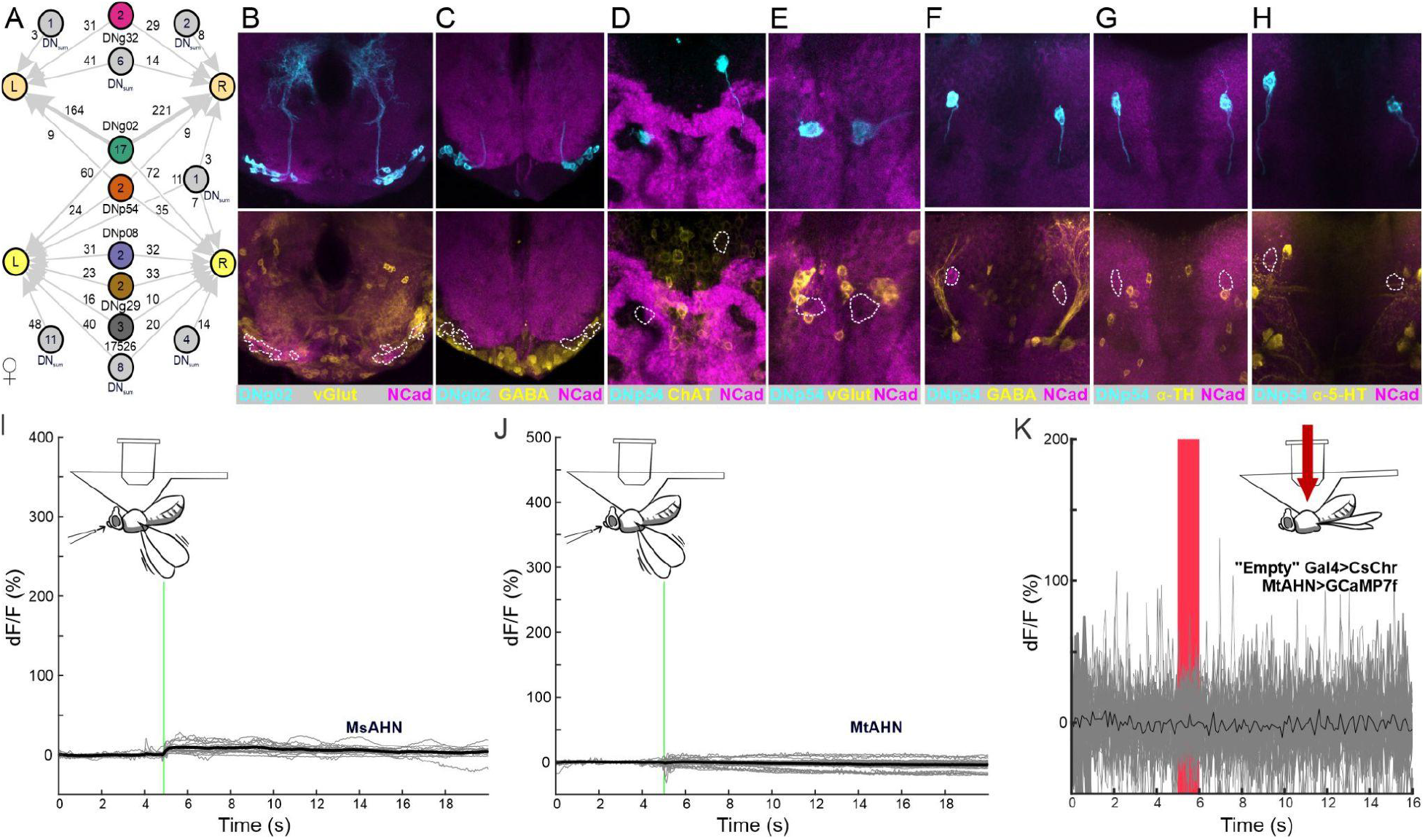
DN input to AHNs in FANC, DN neurotransmitter usage, and AHN calcium imaging controls. **A)** Graph plot of upstream DNs to the MsAHNs and MtAHNs in the FANC dataset. **B)** Intersection between the DNg02 splitGal4 (cyan) and a vGlut-T2A-LexA (yellow) driver line reveals that the DNg02s are not glutamatergic. **C)** The DNg02s (cyan) does not immunolabel for GABA (yellow). **D-H)** The DNp54 splitGal4 line (cyan) does not overlap with T2A-LexAs for **D)** ChAT (yellow) or **E)** vGlut (yellow), nor do the DNp54s immunolabel for **F)** GABA (yellow), **G)** tyrosine hydroxylase (TH; yellow) or **H)** serotonin (5-HT; yellow). NCAD (magenta) delineates neuropil. **I-J**) Flight-induced changes in fluorescence due to movement measured via GFP expression in **I**) the MsAHNs (6 flies, 6 soma, 3 trials) and **J)** the MtAHNs (9 flies, 13 soma, 3 trials). Cartoon depicts orientation of flies during each recording and green line indicates timing of an air puff to trigger flight. Gray traces represent recordings from individual AHN soma and black trace represents the average fluorescence transient across all animals. **K)** Ca^2+^ transients evoked from the MtAHNs in response to CsChrimson activation of an “empty” Gal4 line. Gray traces represent recordings from individual AHN soma and black trace represents the average fluorescence transient across all animals.

**Figure S4.**
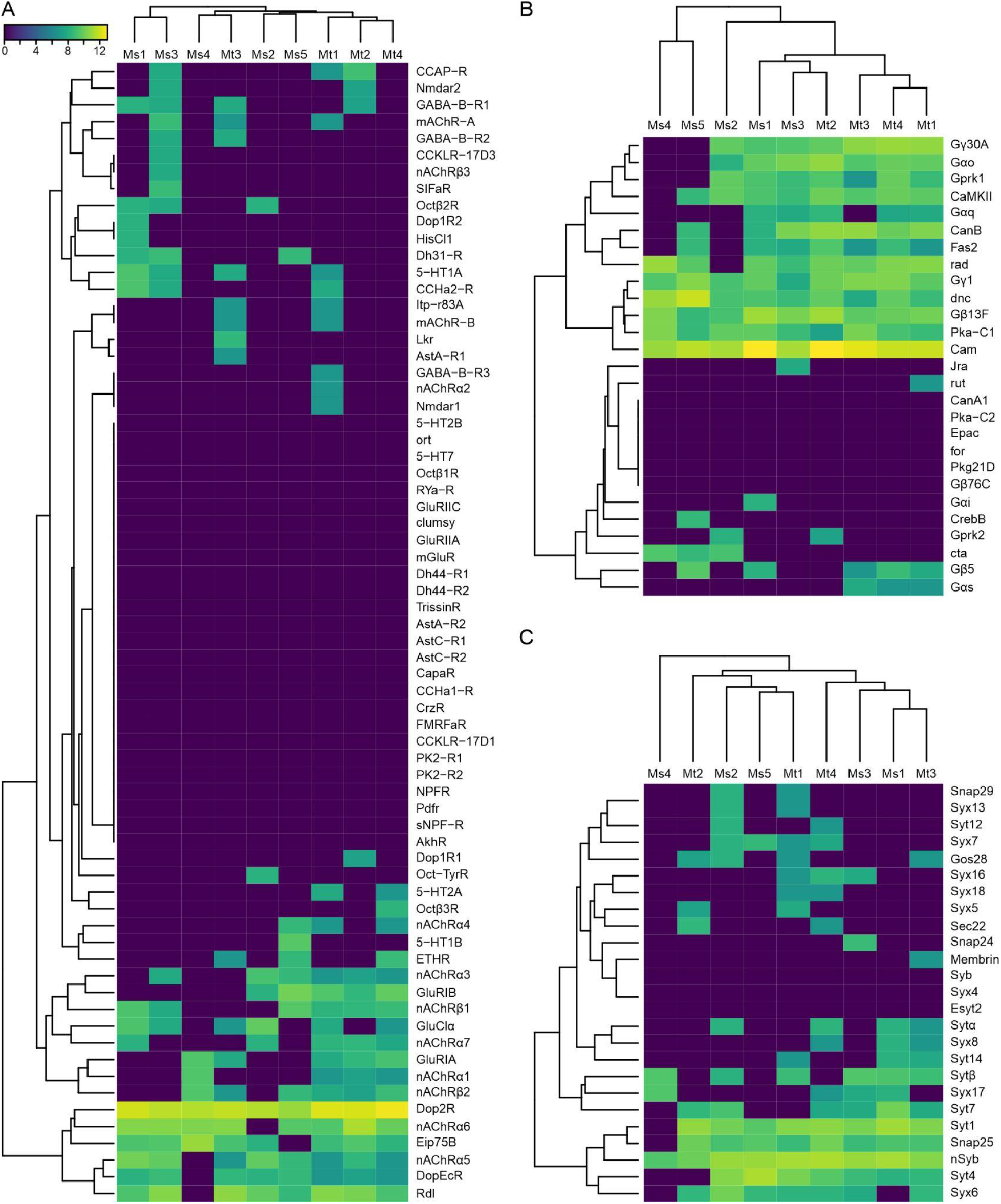
AHN expression profiling. **A)** Hierarchically-clustered heatmap showing normalized expression of neurotransmitter receptor genes screened in cluster analysis. **B)** Hierarchical cluster analysis showing normalized 2nd messenger associated gene product expression across candidate AHNs. **C)** Hierarchical cluster analysis showing normalized expression of synaptic vesicle transmission genes across AHN candidates. Values are read counts of each gene normalized by total counts per million (CPM) per cell, then log scaled (log2(n+1)).

**Figure S5.**
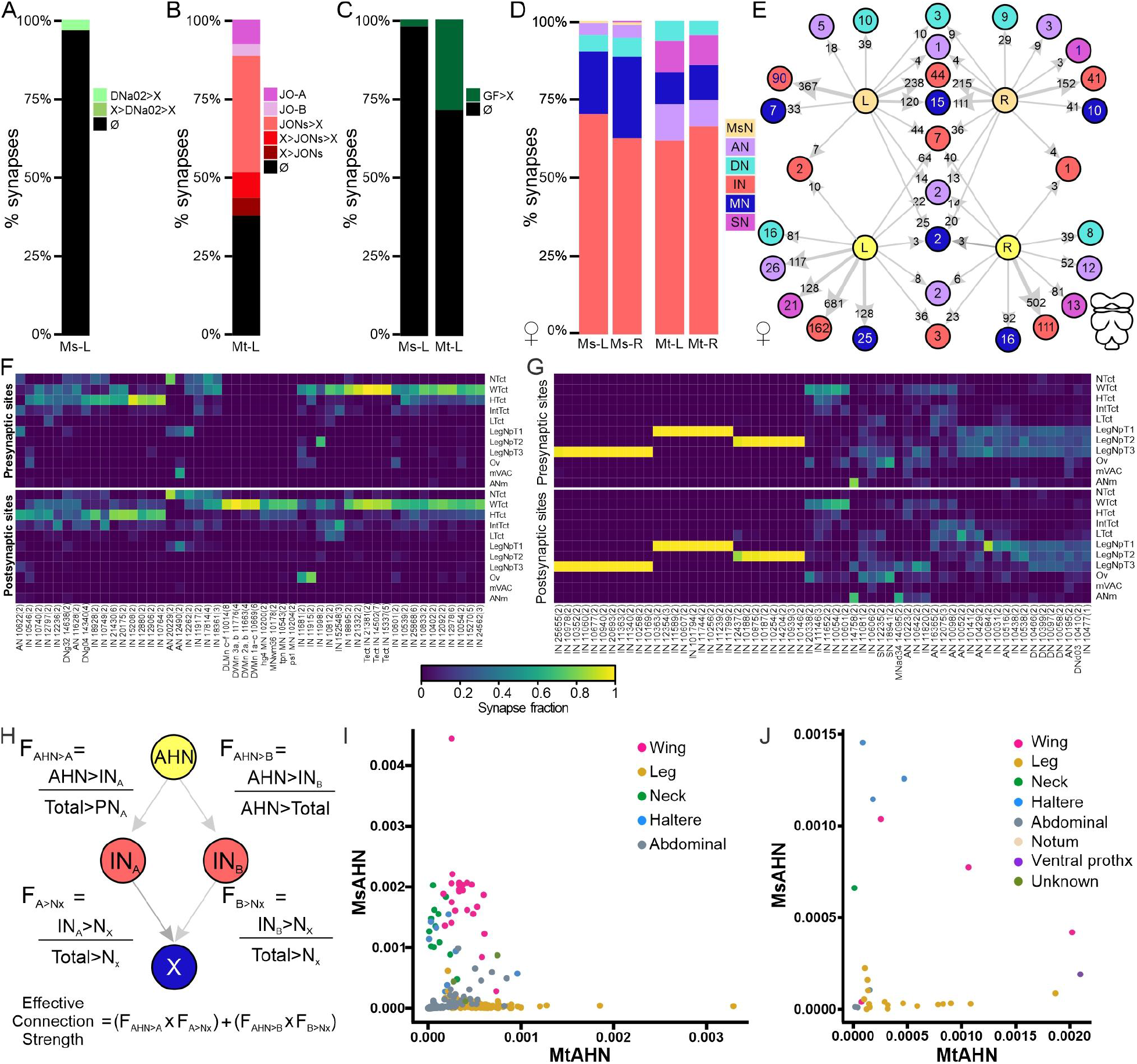
Further information about AHN downstream partners. **A)** Synapse fractions for MsAHN downstream partners in FAFB highlighting those with the DNa02s whether it be upstream (“X>DNa02”) or reciprocal (“X>DNa02>X”) connectivity. “∅fl” indicates neurons with no connectivity to DNa02. **B)** Synapse fraction for the downstream partners of the MtAHN in FAFB classified based on JON type, or connectivity to the JONs otherwise. “X>JONs” indicates neurons upstream of the JONs, “X>JONs>X” indicates neurons both up and downstream of the JONs (threshold of 2 synapses), and “JONs>X” indicates neurons downstream of the JONs. Connectivity of AHN downstream cells with JONs were stochastically traced, and cell count connected to JONs are likely underestimated. **C)** Synapse fractions for MsAHN and MtAHN downstream partners in FAFB that are upstream from from the Giant Fiber neurons (“X>GF”). **D)** Synapse fractions for the downstream partners of the MsAHNs and MtAHNs from the FANC dataset. **E)** Graph plot for the downstream partners of the MsAHNs (peach) and MtAHNs (yellow) from the FANC dataset. **F-G)** Normalized proportion of presynaptic or postsynaptic sites of downstream targets of the **F)** MsAHNs and **G)** MtAHNs plotted based on VNC neuropil within the MANC dataset. Only downstream partners of groups above an outlier threshold of synapse counts with AHNs (above the 3rd quartile plus 1.5 x interquartile range) were included. Neuropil included were based on neck tectulum (NTct), wing tectulum (WTct), haltere tectulum (HTct), intermediate tectulum (IntTct), lower tectulum (LTct), prothoracic leg neuropil (LegNp.T1), mesothoracic leg neuropil (LegNp.T2), metathoracic leg neuropil (LegNp.T3), ovoid/accessory mesothoracic neuropil (Ov), medial ventral association center (mVAC), abdominal neuromeres (ANm). **H)** Cartoon representation of effective connection strength calculations used in Figure 6J and K. F_AHN>A_ is the proportion of the total synapses to a given interneuron (“IN_A_”) that are provided by a given AHN, and F_A>Nx_ is the proportion of the total synapse to a given downstream neuron (“N_x_”) provided by IN_A_. The effective connection strength is therefore the sum of the products of F_AHN>A_ and F_A>Nx_ for every downstream interneuron that is upstream of Nx. **I-J)** Scatterplot of the adjacency scores for all **I)** motor neurons or **j)** sensory neurons within the VNC relative to both AHN pairs. Color-coding is based on body part; wing (pink), leg (gold), neck (green), haltere (blue), abdominal (grey), notum (pink), ventral prothorax and unknown (dark green).

**Figure S6.**
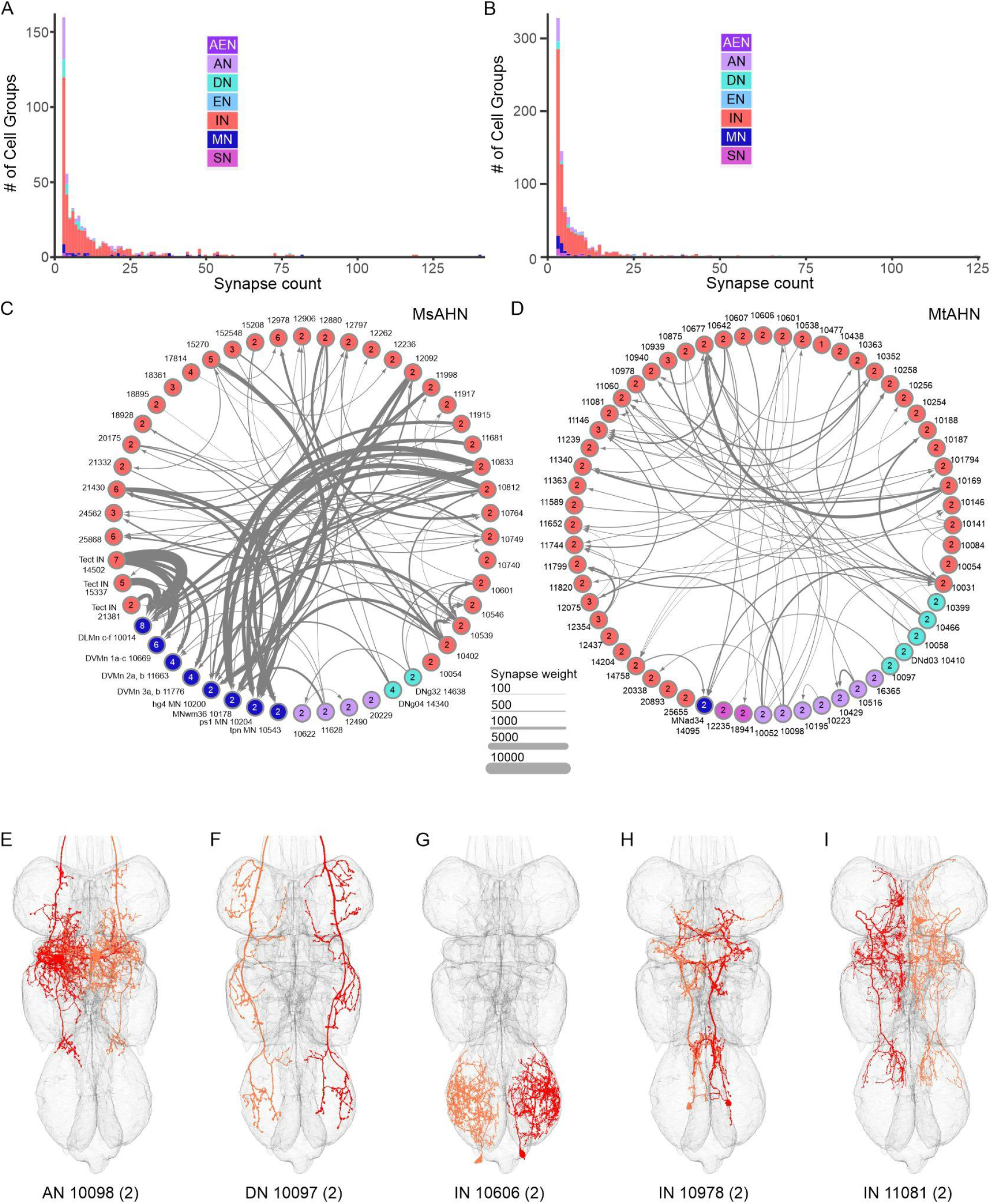
Synapse count distribution, interconnectivity and morphology of AHN downstream partners. **A-B)** Synapse count distribution of **A)** MsAHN and **B)** MtAHN downstream partner groups color coded based on cell-class in MANC. **C-D)** Graph plot depicting the interconnectivity of the downstream partners of **C)** the MsAHNs and **D)** the MtAHNs. Only synapse weights of 100 and above are shown. **E-I)** Reconstructions of the top 5 downstream targets of the MtAHNs in MANC by connectivity strength.

## Acknowledgements

This work was funded by an AFOSR FA9550-17-1-0117, NSF 2127379 to KCD and AMD, a Howard Hughes Medical Institute Janelia Research Campus Visiting Scientist Program project to KCD, AMD, HSJC and GMC, and NIMH117808 to WAL. We thank Maggie Robertson, Emily Tenshaw, Julie Kovalyak, Danielle Matheny, Malia Miller, Emmanuel Mbamalu, Jason Polsky, Shada Alghailani, Melissa Ryan, Srividya Murthy, and Ruchi Parekh for assistance with tracing and proofreading efforts in EM datasets and Danielle Matheny for assistance with collecting confocal optical stacks. We thank Dr. Phillip Chapman for assistance with early experiments in this project. We thank Drs. Tyler Sizemore, Kaylynn Coates, Julius Jonaitis, Katharina Eichler, Gregory Jefferis for technical advice. We thank Drs. Katharina Eichler, Gregory Jefferis and Marta Costa for access to light microscopy level identification of descending neurons in the male adult nerve cord dataset. We thank Dr. Gregory Jefferis for pre-publication access to the malevnc R package. We thank Shigehiro Namiki, the Janelia FlyLight Project team, Janelia Fly Descending Interneuron Project team and Janelia Project Technical Resources for providing pre-publication access to light-level descending neuron typing information including that of DNp54, image of the SS02625 (DNg02) splitGal4 expression pattern, and screening of the DNp54 splitGal4 line SS96091 (VT056359AD; VT002021DBD). Ongoing development of the natverse including the fafbseg package is supported by the NIH BRAIN Initiative (grant 1RF1MH120679-01) and the Medical Research Council (MC_U105188491). MB received support from the Summer Undergraduate Research Experience program at WVU.

## Bibliography

1. von Holst, E., and Mittelstaedt, H. (1950). Das Reafferenzprinzip. Naturwissenschaften 37, 464–476.

2. Straka, H., Simmers, J., and Chagnaud, B.P. (2018). A new perspective on predictive motor signaling. Curr. Biol. 28, R232–R243.

3. Fee, M.S., Mitra, P.P., and Kleinfeld, D. (1997). Central versus peripheral determinants of patterned spike activity in rat vibrissa cortex during whisking. J. Neurophysiol. 78, 1144–1149.

4. Chapman, P.D., Burkland, R., Bradley, S.P., Houot, B., Bullman, V., Dacks, A.M., and Daly, K.C. (2018). Flight motor networks modulate primary olfactory processing in the moth Manduca sexta. Proc Natl Acad Sci USA 115, 5588–5593.

5. Chagnaud, B.P., and Bass, A.H. (2013). Vocal corollary discharge communicates call duration to vertebrate auditory system. J. Neurosci. 33, 18775–18780.

6. Schneider, D.M., Nelson, A., and Mooney, R. (2014). A synaptic and circuit basis for corollary discharge in the auditory cortex. Nature 513, 189–194.

7. Poulet, J.F.A., and Hedwig, B. (2006). The cellular basis of a corollary discharge. Science 311, 518–522.

8. Kim, A.J., Fitzgerald, J.K., and Maimon, G. (2015). Cellular evidence for efference copy in Drosophila visuomotor processing. Nat. Neurosci. 18, 1247–1255.

9. Suga, N., and Jen, P.H. (1975). Peripheral control of acoustic signals in the auditory system of echolocating bats. J. Exp. Biol. 62, 277–311.

10. Drexl, M., and Kössl, M. (2003). Sound-evoked efferent effects on cochlear mechanics of the mustached bat. Hear. Res. 184, 61–74.

11. McGrath, J.J., Saha, S., Al-Hamzawi, A., Alonso, J., Bromet, E.J., Bruffaerts, R., Caldas-de-Almeida, J.M., Chiu, W.T., de Jonge, P., Fayyad, J., et al. (2015). Psychotic Experiences in the General Population: A Cross-National Analysis Based on 31,261 Respondents From 18 Countries. JAMA Psychiatry 72, 697–705.

12. Morris, G.O. (1960). Misperception and disorientation during sleep deprivation. Arch. Gen. Psychiatry 2, 247.

13. Feinberg, I., and Guazzelli, M. (1999). Schizophrenia--a disorder of the corollary discharge systems that integrate the motor systems of thought with the sensory systems of consciousness. Br. J. Psychiatry 174, 196–204.

14. Ford, J.M., and Mathalon, D.H. (2004). Electrophysiological evidence of corollary discharge dysfunction in schizophrenia during talking and thinking. J. Psychiatr. Res. 38, 37–46.

15. Goldberg, R.L., and Henson, O.W. (1998). Changes in cochlear mechanics during vocalization: evidence for a phasic medial efferent effect. Hear. Res. 122, 71–81.

16. Hage, S.R., Jürgens, U., and Ehret, G. (2006). Audio-vocal interaction in the pontine brainstem during self-initiated vocalization in the squirrel monkey. Eur. J. Neurosci. 23, 3297–3308.

17. Eliades, S.J., and Wang, X. (2008). Neural substrates of vocalization feedback monitoring in primate auditory cortex. Nature 453, 1102–1106.

18. Bradley, S.P., Chapman, P.D., Lizbinski, K.M., Daly, K.C., and Dacks, A.M. (2016). A Flight Sensory-Motor to Olfactory Processing Circuit in the Moth Manduca sexta. Front. Neural Circuits 10, 5.

19. Chapman, P.D., Bradley, S.P., Haught, E.J., Riggs, K.E., Haffar, M.M., Daly, K.C., and Dacks, A.M. (2017). Co-option of a motor-to-sensory histaminergic circuit correlates with insect flight biomechanics. Proc. Biol. Sci. 284.

20. Sane, S.P. (2006). Induced airflow in flying insects I. A theoretical model of the induced flow. J. Exp. Biol. 209, 32–42.

21. Sane, S.P., and Jacobson, N.P. (2006). Induced airflow in flying insects II. Measurement of induced flow. J. Exp. Biol. 209, 43–56.

22. Nässel, D.R., Pirvola, U., and Panula, P. (1990). Histaminelike immunoreactive neurons innervating putative neurohaemal areas and central neuropil in the thoraco-abdominal ganglia of the flies Drosophila and Calliphora. J. Comp. Neurol. 297, 525–536.

23. Pollack, I., and Hofbauer, A. (1991). Histamine-like immunoreactivity in the visual system and brain of Drosophila melanogaster. Cell Tissue Res. 266, 391–398.

24. Nässel, D.R., and Elekes, K. (1992). Aminergic neurons in the brain of blowflies and Drosophila: dopamine- and tyrosine hydroxylase-immunoreactive neurons and their relationship with putative histaminergic neurons. Cell Tissue Res. 267, 147–167.

25. Melzig, J., Buchner, S., Wiebel, F., Wolf, R., Burg, M., Pak, W.L., and Buchner, E. (1996). Genetic depletion of histamine from the nervous system of Drosophila eliminates specific visual and mechanosensory behavior. Journal of Comparative Physiology a-Sensory Neural and Behavioral Physiology 179, 763–773.

26. Buchner, E., Buchner, S., Burg, M.G., Hofbauer, A., Pak, W.L., and Pollack, I. (1993). Histamine is a major mechanosensory neurotransmitter candidate in Drosophila melanogaster. Cell Tissue Res. 273, 119–125.

27. Han, D.D., Stein, D., and Stevens, L.M. (2000). Investigating the function of follicular subpopulations during Drosophila oogenesis through hormone-dependent enhancer-targeted cell ablation. Development 127, 573–583.

28. Nern, A., Pfeiffer, B.D., and Rubin, G.M. (2015). Optimized tools for multicolor stochastic labeling reveal diverse stereotyped cell arrangements in the fly visual system. Proc Natl Acad Sci USA 112, E2967–76.

29. Zheng, Z., Lauritzen, J.S., Perlman, E., Robinson, C.G., Nichols, M., Milkie, D., Torrens, O., Price, J., Fisher, C.B., Sharifi, N., et al. (2018). A Complete Electron Microscopy Volume of the Brain of Adult Drosophila melanogaster. Cell 174, 730–743.e22.

30. Phelps, J.S., Hildebrand, D.G.C., Graham, B.J., Kuan, A.T., Thomas, L.A., Nguyen, T.M., Buhmann, J., Azevedo, A.W., Sustar, A., Agrawal, S., et al. (2021). Reconstruction of motor control circuits in adult Drosophila using automated transmission electron microscopy. Cell 184, 759–774.e18.

31. Azevedo, A.W., Lesser, E., Mark, B., Phelps, J.S., Elabbady, L., Kuroda, S., Sustar, A.E., Moussa, A.J., Kandelwal, A., Dallmann, C.J., et al. (2022). Tools for comprehensive reconstruction and analysis of *Drosophila* motor circuits. BioRxiv.

32. Marin, E.C., Morris, B.J., Stuerner, T., Champion, A.S., Krzeminski, D., Badalamente, G., Gkantia, M., Dunne, C.R., Eichler, K., Takemura, S., et al. (2023). Systematic annotation of a complete adult male Drosophila nerve cord connectome reveals principles of functional organisation. BioRxiv.

33. Takemura, S., Hayworth, K.J., Huang, G.B., Januszewski, M., Lu, Z., Marin, E.C., Preibisch, S., Xu, C.S., Bogovic, J., Champion, A.S., et al. (2023). A connectome of the male *drosophila* ventral nerve cord. BioRxiv.

34. Costa, M., Manton, J.D., Ostrovsky, A.D., Prohaska, S., and Jefferis, G.S.X.E. (2016). NBLAST: rapid, sensitive comparison of neuronal structure and construction of neuron family databases. Neuron 91, 293–311.

35. Nicolaï, L.J.J., Ramaekers, A., Raemaekers, T., Drozdzecki, A., Mauss, A.S., Yan, J., Landgraf, M., Annaert, W., and Hassan, B.A. (2010). Genetically encoded dendritic marker sheds light on neuronal connectivity in Drosophila. Proc Natl Acad Sci USA 107, 20553–20558.

36. Namiki, S., Dickinson, M.H., Wong, A.M., Korff, W., and Card, G.M. (2018). The functional organization of descending sensory-motor pathways in Drosophila. eLife 7.

37. Court, R., Namiki, S., Armstrong, J.D., Börner, J., Card, G., Costa, M., Dickinson, M., Duch, C., Korff, W., Mann, R., et al. (2020). A systematic nomenclature for the drosophila ventral nerve cord. Neuron 107, 1071–1079.e2.

38. Namiki, S., Ros, I.G., Morrow, C., Rowell, W.J., Card, G.M., Korff, W., and Dickinson, M.H. (2022). A population of descending neurons that regulates the flight motor of Drosophila. Curr. Biol. 32, 1189–1196.e6.

39. Cheong, H.S.J., Eichler, K., Stuerner, T., Asinof, S.K., Champion, A.S., Marin, E.C., Oram, T.B., Sumathipala, M., Venkatasubramanian, L., Namiki, S., et al. (2023). Transforming descending input into behavior: The organization of premotor circuits in the Drosophila Male Adult Nerve Cord connectome | bioRxiv. Available at: https://www.biorxiv.org/content/10.1101/2023.06.07.543976v1 [Accessed June 6, 2023].

40. Palmer, E.H., Omoto, J.J., and Dickinson, M.H. (2022). The role of a population of descending neurons in the optomotor response in flying Drosophila. BioRxiv.

41. Diao, F., Ironfield, H., Luan, H., Diao, F., Shropshire, W.C., Ewer, J., Marr, E., Potter, C.J., Landgraf, M., and White, B.H. (2015). Plug-and-play genetic access to drosophila cell types using exchangeable exon cassettes. Cell Rep. 10, 1410–1421.

42. Klapoetke, N.C., Murata, Y., Kim, S.S., Pulver, S.R., Birdsey-Benson, A., Cho, Y.K., Morimoto, T.K., Chuong, A.S., Carpenter, E.J., Tian, Z., et al. (2014). Independent optical excitation of distinct neural populations. Nat. Methods 11, 338–346.

43. Allen, A.M., Neville, M.C., Birtles, S., Croset, V., Treiber, C.D., Waddell, S., and Goodwin, S.F. (2020). A single-cell transcriptomic atlas of the adult Drosophila ventral nerve cord. eLife 9.

44. Lannutti, B.J., and Schneider, L.E. (2001). Gprk2 controls cAMP levels in Drosophila development. Dev. Biol. 233, 174–185.

45. Nighorn, A., Healy, M.J., and Davis, R.L. (1991). The cyclic AMP phosphodiesterase encoded by the Drosophila dunce gene is concentrated in the mushroom body neuropil. Neuron 6, 455–467.

46. Silies, M., Gohl, D.M., and Clandinin, T.R. (2014). Motion-detecting circuits in flies: coming into view. Annu. Rev. Neurosci. 37, 307–327.

47. Zhu, Y. (2013). The Drosophila visual system: From neural circuits to behavior. Cell Adh. Migr. 7, 333–344.

48. Borst, A., and Haag, J. (2002). Neural networks in the cockpit of the fly. J. Comp. Physiol. A Neuroethol. Sens. Neural Behav. Physiol. 188, 419–437.

49. Levine, J., and Tracey, D. (1973). Structure and function of the giant motorneuron ofDrosophila melanogaster. J. Comp. Physiol. 87, 213–235.

50. Bacon, J.P., and Strausfeld, N.J. (1986). The dipteran ?Giant fibre? pathway: neurons and signals. J. Comp. Physiol. 158, 529–548.

51. Mu, L., Bacon, J.P., Ito, K., and Strausfeld, N.J. (2014). Responses of Drosophila giant descending neurons to visual and mechanical stimuli. J. Exp. Biol. 217, 2121–2129.

52. von Reyn, C.R., Nern, A., Williamson, W.R., Breads, P., Wu, M., Namiki, S., and Card, G.M. (2017). Feature integration drives probabilistic behavior in the drosophila escape response. Neuron 94, 1190–1204.e6.

53. Ache, J.M., Polsky, J., Alghailani, S., Parekh, R., Breads, P., Peek, M.Y., Bock, D.D., von Reyn, C.R., and Card, G.M. (2019). Neural basis for looming size and velocity encoding in the drosophila giant fiber escape pathway. Curr. Biol. 29, 1073–1081.e4.

54. Rayshubskiy, A., Holtz, S.L., D’Alessandro, I., Li, A.A., Vanderbeck, Q.X., Haber, I.S., Gibb, P.W., and Wilson, R.I. (2020). Neural circuit mechanisms for steering control in walking *Drosophila*. BioRxiv.

55. Strausfeld, N.J., and Bassemir, U.K. (1985). Lobula plate and ocellar interneurons converge onto a cluster of descending neurons leading to neck and leg motor neuropil in Calliphora erythrocephala. Cell Tissue Res. 240, 617–640.

56. Namiki, S., and Kanzaki, R. (2016). Comparative neuroanatomy of the lateral accessory lobe in the insect brain. Front. Physiol. 7, 244.

56. Boergens, K.M., Kapfer, C., Helmstaedter, M., Denk, W., and Borst, A. (2018). Full reconstruction of large lobula plate tangential cells in Drosophila from a 3D EM dataset. PLoS ONE 13, e0207828.

58. Yorozu, S., Wong, A., Fischer, B.J., Dankert, H., Kernan, M.J., Kamikouchi, A., Ito, K., and Anderson, D.J. (2009). Distinct sensory representations of wind and near-field sound in the Drosophila brain. Nature 458, 201–205.

59. Ishikawa, Y., and Kamikouchi, A. (2016). Auditory system of fruit flies. Hear. Res. 338, 1–8.

60. Todi, S.V., Sharma, Y., and Eberl, D.F. (2004). Anatomical and molecular design of the Drosophila antenna as a flagellar auditory organ. Microsc. Res. Tech. 63, 388–399.

61. Caldwell, J.C., and Eberl, D.F. (2002). Towards a molecular understanding of Drosophila hearing. J. Neurobiol. 53, 172–189.

62. Matsuo, E., and Kamikouchi, A. (2013). Neuronal encoding of sound, gravity, and wind in the fruit fly. J. Comp. Physiol. A Neuroethol. Sens. Neural Behav. Physiol. 199, 253–262.

63. Lai, J.S.-Y., Lo, S.-J., Dickson, B.J., and Chiang, A.-S. (2012). Auditory circuit in the Drosophila brain. Proc Natl Acad Sci USA 109, 2607–2612.

64. Tootoonian, S., Coen, P., Kawai, R., and Murthy, M. (2012). Neural representations of courtship song in the Drosophila brain. J. Neurosci. 32, 787–798.

65. Matsuo, E., Seki, H., Asai, T., Morimoto, T., Miyakawa, H., Ito, K., and Kamikouchi, A. (2016). Organization of projection neurons and local neurons of the primary auditory center in the fruit fly Drosophila melanogaster. J. Comp. Neurol. 524, 1099–1164.

66. Matsuo, E., Yamada, D., Ishikawa, Y., Asai, T., Ishimoto, H., and Kamikouchi, A. (2014). Identification of novel vibration- and deflection-sensitive neuronal subgroups in Johnston’s organ of the fruit fly. Front. Physiol. 5, 179.

67. Vaughan, A.G., Zhou, C., Manoli, D.S., and Baker, B.S. (2014). Neural pathways for the detection and discrimination of conspecific song in D. melanogaster. Curr. Biol. 24, 1039–1049.

68. Patella, P., and Wilson, R.I. (2018). Functional maps of mechanosensory features in the drosophila brain. Curr. Biol. 28, 1189–1203.e5.

69. Pézier, A., Jezzini, S.H., Marie, B., and Blagburn, J.M. (2014). Engrailed alters the specificity of synaptic connections of Drosophila auditory neurons with the giant fiber. J. Neurosci. 34, 11691–11704.

70. Lehnert, B.P., Baker, A.E., Gaudry, Q., Chiang, A.-S., and Wilson, R.I. (2013). Distinct roles of TRP channels in auditory transduction and amplification in Drosophila. Neuron 77, 115–128.

71. Kim, H., Horigome, M., Ishikawa, Y., Li, F., Lauritzen, J.S., Card, G., Bock, D.D., and Kamikouchi, A. (2020). Wiring patterns from auditory sensory neurons to the escape and song-relay pathways in fruit flies. J. Comp. Neurol. 528, 2068–2098.

72. Li, F., Lindsey, J.W., Marin, E.C., Otto, N., Dreher, M., Dempsey, G., Stark, I., Bates, A.S., Pleijzier, M.W., Schlegel, P., et al. (2020). The connectome of the adult Drosophila mushroom body provides insights into function. eLife 9.

73. Buchanan, J.T., and Einum, J.F. (2008). The spinobulbar system in lamprey. Brain Res. Rev. 57, 37–45.

74. Stecina, K., Fedirchuk, B., and Hultborn, H. (2013). Information to cerebellum on spinal motor networks mediated by the dorsal spinocerebellar tract. J Physiol (Lond) 591, 5433–5443.

75. Chagnaud, B.P., Banchi, R., Simmers, J., and Straka, H. (2015). Spinal corollary discharge modulates motion sensing during vertebrate locomotion. Nat. Commun. 6, 7982.

76. Crapse, T.B., and Sommer, M.A. (2008). Corollary discharge across the animal kingdom. Nat. Rev. Neurosci. 9, 587–600.

77. Fukutomi, M., and Carlson, B.A. (2020). A history of corollary discharge: contributions of mormyrid weakly electric fish. Front. Integr. Neurosci. 14, 42.

78. Maimon, G., Straw, A.D., and Dickinson, M.H. (2010). Active flight increases the gain of visual motion processing in Drosophila. Nat. Neurosci. 13, 393–399.

79. Suver, M.P., Mamiya, A., and Dickinson, M.H. (2012). Octopamine neurons mediate flight-induced modulation of visual processing in Drosophila. Curr. Biol. 22, 2294–2302.

80. Jung, S.N., Borst, A., and Haag, J. (2011). Flight activity alters velocity tuning of fly motion-sensitive neurons. J. Neurosci. 31, 9231–9237.

81. van Breugel, F., Suver, M.P., and Dickinson, M.H. (2014). Octopaminergic modulation of the visual flight speed regulator of Drosophila. J. Exp. Biol. 217, 1737–1744.

82. Ache, J.M., Namiki, S., Lee, A., Branson, K., and Card, G.M. (2019). State-dependent decoupling of sensory and motor circuits underlies behavioral flexibility in Drosophila. Nat. Neurosci. 22, 1132–1139.

83. Stuart, G.J., and Spruston, N. (2015). Dendritic integration: 60 years of progress. Nat. Neurosci. 18, 1713–1721.

84. Cover, K.K., and Mathur, B.N. (2021). Axo-axonic synapses: Diversity in neural circuit function. J. Comp. Neurol. 529, 2391–2401.

85. Poulet, J.F.A., and Hedwig, B. (2002). A corollary discharge maintains auditory sensitivity during sound production. Nature 418, 872–876.

86. Poulet, J.F.A., and Hedwig, B. (2003). Corollary discharge inhibition of ascending auditory neurons in the stridulating cricket. J. Neurosci. 23, 4717–4725.

87. Bidaye, S.S., Laturney, M., Chang, A.K., Liu, Y., Bockemühl, T., Büschges, A., and Scott, K. (2020). Two brain pathways initiate distinct forward walking programs in drosophila. Neuron 108, 469–485.e8.

88. Fujiwara, T., Brotas, M., and Chiappe, M.E. (2022). Walking strides direct rapid and flexible recruitment of visual circuits for course control in Drosophila. Neuron 110, 2124–2138.e8.

89. Feng, K., Palfreyman, M.T., Häsemeyer, M., Talsma, A., and Dickson, B.J. (2014). Ascending SAG neurons control sexual receptivity of Drosophila females. Neuron 83, 135–148.

90. Mann, K., Gordon, M.D., and Scott, K. (2013). A pair of interneurons influences the choice between feeding and locomotion in Drosophila. Neuron 79, 754–765.

91. Kim, H., Kirkhart, C., and Scott, K. (2017). Long-range projection neurons in the taste circuit of Drosophila. eLife 6.

92. Tuthill, J.C., and Wilson, R.I. (2016). Parallel transformation of tactile signals in central circuits of drosophila. Cell 164, 1046–1059.

93. Sen, R., Wang, K., and Dickson, B.J. (2019). TwoLumps Ascending Neurons Mediate Touch-Evoked Reversal of Walking Direction in Drosophila. Curr. Biol. 29, 4337–4344.e5.

94. Bidaye, S.S., Machacek, C., Wu, Y., and Dickson, B.J. (2014). Neuronal control of Drosophila walking direction. Science 344, 97–101.

95. Hampel, S., McKellar, C.E., Simpson, J.H., and Seeds, A.M. (2017). Simultaneous activation of parallel sensory pathways promotes a grooming sequence in Drosophila. eLife 6.

96. Chen, C.-L., Aymanns, F., Minegishi, R., Matsuda, V., Talabot, N., Gunel, S., Dickson, B.J., and Ramdya, P. (2022). Ascending neurons convey behavioral state to integrative sensory and action selection centers in the brain. BioRxiv.

97. Jenett, A., Rubin, G.M., Ngo, T.-T.B., Shepherd, D., Murphy, C., Dionne, H., Pfeiffer, B.D., Cavallaro, A., Hall, D., Jeter, J., et al. (2012). A GAL4-driver line resource for Drosophila neurobiology. Cell Rep. 2, 991–1001.

98. Pfeiffer, B.D., Ngo, T.-T.B., Hibbard, K.L., Murphy, C., Jenett, A., Truman, J.W., and Rubin, G.M. (2010). Refinement of tools for targeted gene expression in Drosophila. Genetics 186, 735–755.

99. Pfeiffer, B.D., Truman, J.W., and Rubin, G.M. (2012). Using translational enhancers to increase transgene expression in Drosophila. Proc Natl Acad Sci USA 109, 6626–6631.

100. Dionne, H., Hibbard, K.L., Cavallaro, A., Kao, J.-C., and Rubin, G.M. (2018). Genetic Reagents for Making Split-GAL4 Lines in Drosophila. Genetics 209, 31–35.

101. Tirian, L., and Dickson, B. (2017). The VT GAL4, LexA, and split-GAL4 driver line collections for targeted expression in the Drosophila nervous system. BioRxiv.

102. Dana, H., Sun, Y., Mohar, B., Hulse, B.K., Kerlin, A.M., Hasseman, J.P., Tsegaye, G., Tsang, A., Wong, A., Patel, R., et al. (2019). High-performance calcium sensors for imaging activity in neuronal populations and microcompartments. Nat. Methods 16, 649–657.

103. Zhang, B., Freeman, M.R., and Waddell, S. (2010). Drosophila neurobiology : a laboratory manual (Cold Spring Harbor, N.Y.: Cold Spring Harbor Laboratory Press).

104. Saalfeld, S., Cardona, A., Hartenstein, V., and Tomancak, P. (2009). CATMAID: collaborative annotation toolkit for massive amounts of image data. Bioinformatics 25, 1984–1986.

105. Schneider-Mizell, C.M., Gerhard, S., Longair, M., Kazimiers, T., Li, F., Zwart, M.F., Champion, A., Midgley, F.M., Fetter, R.D., Saalfeld, S., et al. (2016). Quantitative neuroanatomy for connectomics in Drosophila. eLife 5.

106. Li, P.H., Lindsey, L.F., Januszewski, M., Zheng, Z., Bates, A.S., Taisz, I., Tyka, M., Nichols, M., Li, F., Perlman, E., et al. (2019). Automated Reconstruction of a Serial-Section EM *Drosophila* Brain with Flood-Filling Networks and Local Realignment. BioRxiv.

107. Bates, A.S., Manton, J.D., Jagannathan, S.R., Costa, M., Schlegel, P., Rohlfing, T., and Jefferis, G.S. (2020). The natverse, a versatile toolbox for combining and analysing neuroanatomical data. eLife 9.

108. Turner-Evans, D., Wegener, S., Rouault, H., Franconville, R., Wolff, T., Seelig, J.D., Druckmann, S., and Jayaraman, V. (2017). Angular velocity integration in a fly heading circuit. eLife 6.

109. Yaksi, E., and Wilson, R.I. (2010). Electrical coupling between olfactory glomeruli. Neuron 67, 1034–1047.

110. Edelstein, A.D., Tsuchida, M.A., Amodaj, N., Pinkard, H., Vale, R.D., and Stuurman, N. (2014). Advanced methods of microscope control using μManager software. J. Biol. Methods 1.

111. Pologruto, T.A., Sabatini, B.L., and Svoboda, K. (2003). ScanImage: flexible software for operating laser scanning microscopes. Biomed. Eng. Online 2, 13.

112. Schindelin, J., Arganda-Carreras, I., Frise, E., Kaynig, V., Longair, M., Pietzsch, T., Preibisch, S., Rueden, C., Saalfeld, S., Schmid, B., et al. (2012). Fiji: an open-source platform for biological-image analysis. Nat. Methods *9*, 676–682.

113. Dubbs, A., Guevara, J., and Yuste, R. (2016). moco: Fast Motion Correction for Calcium Imaging. Front. Neuroinformatics 10, 6.

